# Ephrin inhibition disrupts stromal-cancer crosstalk and reduces metastasis in pancreatic cancer

**DOI:** 10.64898/2025.12.10.693406

**Authors:** Cerian L. Bolton, Rhianna O’Sullivan, Guillem Fuertes-Marin, Francis Sprouse, Hanghang Wang, Shinelle Menezes, Amandine C. Tan, Sabrina Simoncelli, Angus J. M. Cameron, John F. Marshall, Mark Henkemeyer, Edward P. Carter, Richard P. Grose

## Abstract

Cancer associated fibroblasts (CAFs) are critical drivers of disease progression and metastasis within the pancreatic tumour microenvironment. Using a 3D spheroid model of CAF-led invasion, we identified complementary expression of ephrin family receptors (EPHB2) and ligands (EPHRINB2) between cancer cells and CAFs, implicating this bidirectional signalling family in tumour progression. Through pancreatic stellate cell-derived CAFs isolated from genetically modified mouse models, where EphrinB was either lacking or modified by a gain-of-function mutation, we identified both forward and reverse signalling to be required for invasion. In a syngeneic murine orthotopic model of pancreatic cancer, we show that first-in-class tetramerisation inhibitors of Eph/Ephrin interactions can reduce local invasion in primary tumours, and significantly decrease metastatic tumour spread across multiple organ sites. Our data highlight Ephrin signalling as a critical nexus for CAF-cancer crosstalk and establish a foundation for the clinical development of targeted EPH-EPHRIN inhibitors to counter metastatic invasion.

## Main

Despite a rising incidence of pancreatic cancer, survival rates continue to be dire, ranking 6^th^ highest for worldwide mortality^1^. Accounting for 90% of cases, pancreatic ductal adenocarcinoma (PDAC) has limited treatment options, with palliative care providing the most common strategy^2, 3^. Delayed diagnosis and high disease severity are underpinned by the desmoplastic nature of the disease^4^, with abundant cancer associated fibroblasts (CAF) driving disease progression and immune evasion^5^. A central role for CAFs is in driving cancer cell invasion, where CAFs can use extracellular proteases to burrow through the extracellular matrix (ECM) and guide cancer cells via cell-cell contacts^6^.

Previous studies have pointed to a key role for receptor tyrosine kinases (RTKs) in PDAC-CAF biology^7–10^. However, research surrounding the largest RTK superfamily, erythropoietin-producing hepatocellular carcinoma receptor (EPH) is limited in PDAC, despite compelling evidence for driving cellular invasion in other contexts^11–15^.

EPH receptors are divided into A and B classes (EPHA1-8, A10 and EPHB1-4, B6), with class separation based upon the EPH protein sequence and corresponding ligand subtype^16^. EPHRIN (EFNA1-5, EFNB1-3) ligands are classed dependent on structural differences, with EPHRINA proteins attached to the cell surface via a GPI-linkage and EPHRINB by a hydrophobic transmembrane domain, followed by a short intracellular tail which contains tyrosine residues and a terminal PDZ domain^16^. Generally, EPHA receptors will bind EPHRINA ligands, and EPHB receptors will bind EPHRINB ligands^16^. Upon receptor-ligand binding, signalling is triggered in a bidirectional manner through both the receptor (forward signalling) and ligand (reverse signalling) ^17^. After dimerisation and autophosphorylation, forward EPH signalling activates common RTK downstream pathways^17^. Reverse EPHRIN signalling dynamics are not as well understood. EPHRINA ligands are known to signal intracellularly as part of lipid rafts, whilst EPHRINB ligands rely upon phosphorylation of intracellular tyrosine residues to recruit SH2/SH3 domain proteins, and PDZ domain containing proteins to the EPHRINB PDZ motif^17^. Full activation of bidirectional signalling requires formation of higher-order tetramers and tetramer superclusters, where signal activity is proportional to cluster size^17^.

Developmentally, EPH signalling is paramount to direct cell migration in axon guidance and angiogenesis, although roles in cell proliferation, death and differentiation are also reported^16^. In cancer, EPH signalling has been associated with both pro- and anti-tumoural effects, with expression increased in a range of cancers, both in cancer cells and within the tumour microenvironment (TME) ^18–20^. For example, EPHB4-expressing ovarian cancer cells and EPHRINB2-expressing endothelial cells have highlighted opposing functions. EPHB4 forward signalling in cancer cells suppressed proliferation, whilst reverse EPHRINB2 signalling in endothelial cells promoted tumour invasion and angiogenesis^13^. This same axis has been reported in head and neck cancer^14^. Interestingly, EPHRINB2-Fc was also found to activate EPHB4 (and EPHB3) in PC-3 prostate cancer cell migration, where knockdown of both receptors restored contact inhibition^15^. In models of pancreatic cancer, EPHRINB2 knockdown could reduce invasion of cancer cells through inhibition of epithelial-mesenchymal transition (EMT) ^11^. The majority of these studies focus predominantly on the role of EPHRIN ligands to initiate forward EPH signalling in cancer, with less exploration of reverse signalling. Limited literature surrounds the role of EPHRIN reverse signalling in PDAC, with our work being the first to address the function of EPHRIN reverse signalling in CAF-PDAC crosstalk.

We hypothesise that EPHRINB2 reverse signalling drives CAF-led invasion of cancer cells. Here, we use genetic and pharmacological approaches, both *in vitro* and *in vivo*, to demonstrate the critical role of EPHRINB2 in CAF invasion, promoting PDAC invasion both through forward EPHB2 signalling in cancer cells and reverse EPHRINB2 signalling in CAFs. Blockade of this bi-directional signalling axis, using a first-in-class tetramerisation inhibitor^21, 22^, results in significantly decreased invasion and metastasis, providing a clear rationale for clinical development of these compounds.

## Results

### A cancer cell – CAF EPHB2 – EPHRINB2 signalling axis drives invasion

We first explored reciprocal communication between CAFs and PDAC cancer cells required for invasion using a multicellular, matrix-embedded, spheroid model (**Figure 1A**). Chimeric heterocellular spheroids containing human cancer cells and mouse CAFs (and *vice versa*) were processed for bulk RNA sequencing (RNA-seq), enabling bioinformatic deconvolution of cell-type specific transcripts based upon species homology. This revealed complementary expression of EPHRIN ligands and EPH receptors between CAFs and cancer cells respectively (**Figure 1B**) ^23^.

**Figure 1.**
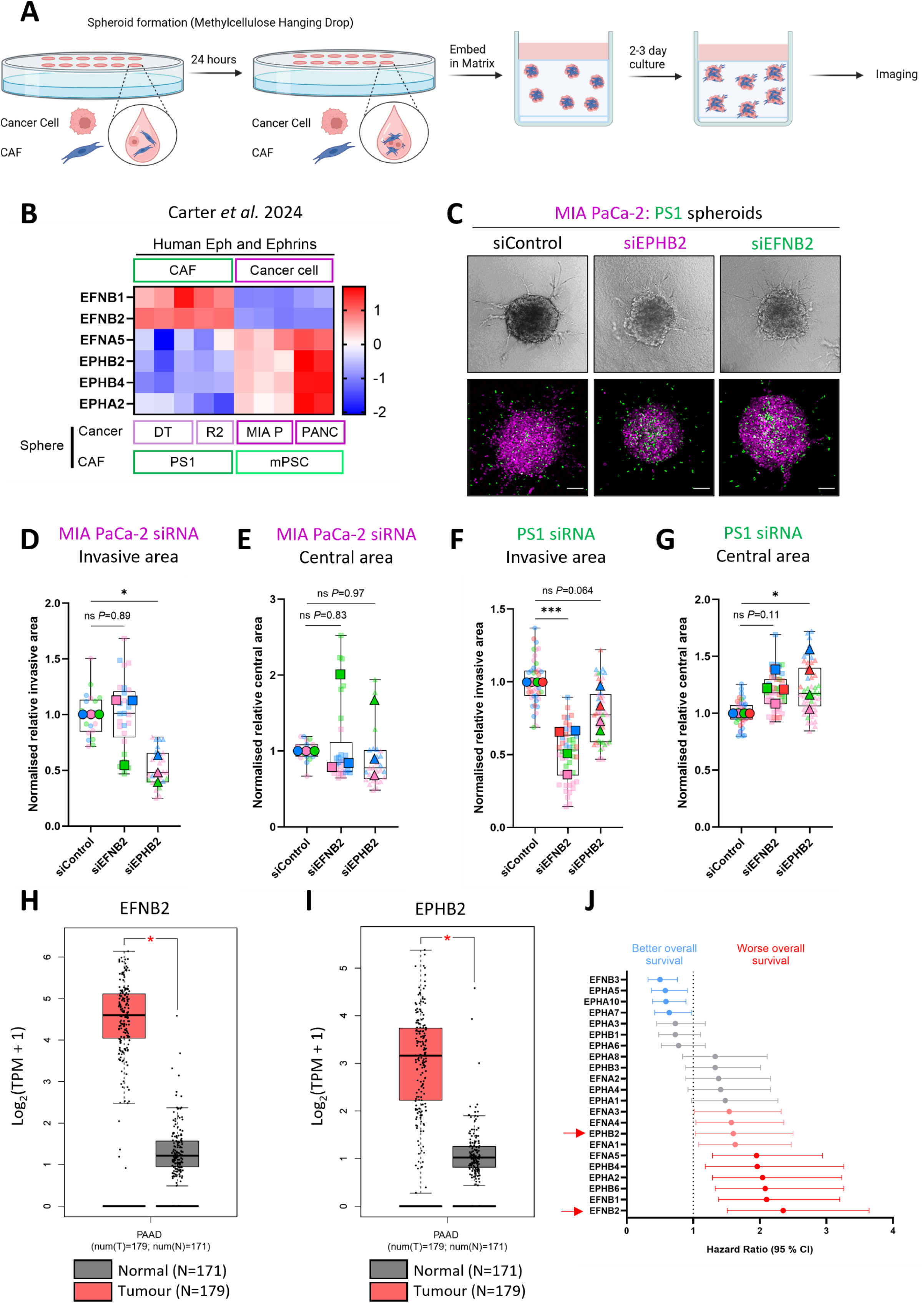
A cancer cell – CAF EPHB2 – EphrinB2 signalling axis drives invasion. (A) Schematic of hanging drop spheroid model. Cancer cells and CAFs form spheres within methylcellulose hanging droplets, which are subsequently embedded into collagen/Matrigel mix and cultured for up to 3 days prior to imaging. (B) Heatmap comparison of EPHRIN ligand and EPH receptor expression in human CAFs and human PDAC cells. Data gained from RNA-seq of chimeric heterocellular spheroids as described in Carter *et al*. 2024^23^. (C) Representative brightfield and confocal images of day 3 chimeric spheroids containing H2B-RFP MIA PaCa-2 human PDAC cells (purple) and H2B-GFP PS1 human CAF cells (green), with cell type specific knockdown of either EPHB2 or EPHRINB2. Scale bar = 100 μm. (D-G) Quantification of MIA PaCa-2/PS1 spheroid invasive (D, F) and central (E, G) area normalised to siRNA control. Graphs presented as superplots (1 point = 1 spheroid), with individual colours representing each biological repeat and averages per biological repeat shown outlined and in bold. Ordinary one-way ANOVA with Dunnett’s multiple comparisons test \**P*<0.05, \*\*\**P*<0.001 (N=3). (H, I) *EFNB2* (H) and *EPHB2* (I) expression in tumour bulk (red) vs. normal tissue (grey). Data were taken from GEPIA using matched pancreatic cancer patient TCGA (PAAD) and GTEx data (Normal N=171, Tumour N=179), Log_2_(fold change) cut off = 1, \**P<*0.01. (J) Hazard ratio (HR) values for each EPHRIN ligand and EPH receptor showing the relationship between overexpression and patient overall survival (blue indicates significantly better overall survival (HR<1), red indicates significantly worse overall survival (HR>1), grey indicates points which were not significant P>0.05). Bold points show the HR value, with error bars indicating HR spread using RNA-seq mRNA data from 177 pancreatic cancer patients. Red arrows point to EPHB2 and EPHRINB2.

To determine whether EPH-EPHRIN crosstalk was involved in invasion, relevant hits were targeted by siRNA knockdown in either MIA PaCa-2 PDAC cells or PS1 CAFs, and invasion evaluated through our spheroid model. A significant decrease in invasion was observed when either EPHB2 was knocked down in cancer cells, or when EPHRINB2 was silenced in CAFs, highlighting this ligand-receptor pair as a potential key axis in regulating invasion (**Figure 1C-G**; **Supplementary Figure 1**). No decreases in central spheroid area were recorded for either cell type when EPHB2 or EPHRINB2 were targeted, indicating knockdown specifically affected invasive potential over cell viability (**Figure 1E, G**).

To compare our observations to clinical PDAC, we performed bioinformatic analysis of publicly available transcriptomic data from patients with pancreatic cancers, identifying significantly increased expression of both *EFNB2* and *EPHB2* in tumour bulk compared to normal tissue (**Figure 1H, I**). Interestingly, high expression of *EFNB2* in pancreatic cancer also correlates with the worst overall survival of any *EPH* or *EFN* (**Figure 1J**). Taken together, these data suggest that EPHB2-EPHRINB2 signalling may be a key axis in CAF-PDAC crosstalk, critical for disease progression.

### EphrinB reverse signalling is required for stromal-led invasion

To determine the importance of EphrinB2 reverse signalling for cellular invasion, we isolated stellate cells (a CAF precursor^24^) from the pancreata of a panel of genetically engineered mice. The panel comprised an *efnb^WT^* line (overexpressing normal EphrinB wild-type (WT)) and an *efnb1/2^-/-^* genetic knockout (**Figure 2A**). We also generated an EphrinB gain-of-function (GOF) line (*efnb^GOF^*). This line exhibits constitutive reverse signalling through a GOF insert, where the intracellular tyrosine residues of EphrinB are replaced with SH3 domains from the EphrinB downstream adaptor Grb4/NCK2/Nckβ^25, 26^ (**Figure 2A**). This GOF bypasses the need for EPH receptor engagement to activate EphrinB reverse signalling, as the attachment of Grb4 SH3 domains will constitutively recruit PXXP proline-rich target cytoskeletal regulatory proteins to the plasma membrane. This is opposed to the normal requirement for EPH engagement of EphrinB to induce tyrosine phosphorylation of the EphrinB intracellular tail, which then leads to recruitment of the Grb4 SH2 domain and its associated SH3-bound cytoskeletal regulatory proteins to the plasma membrane.

**Figure 2.**
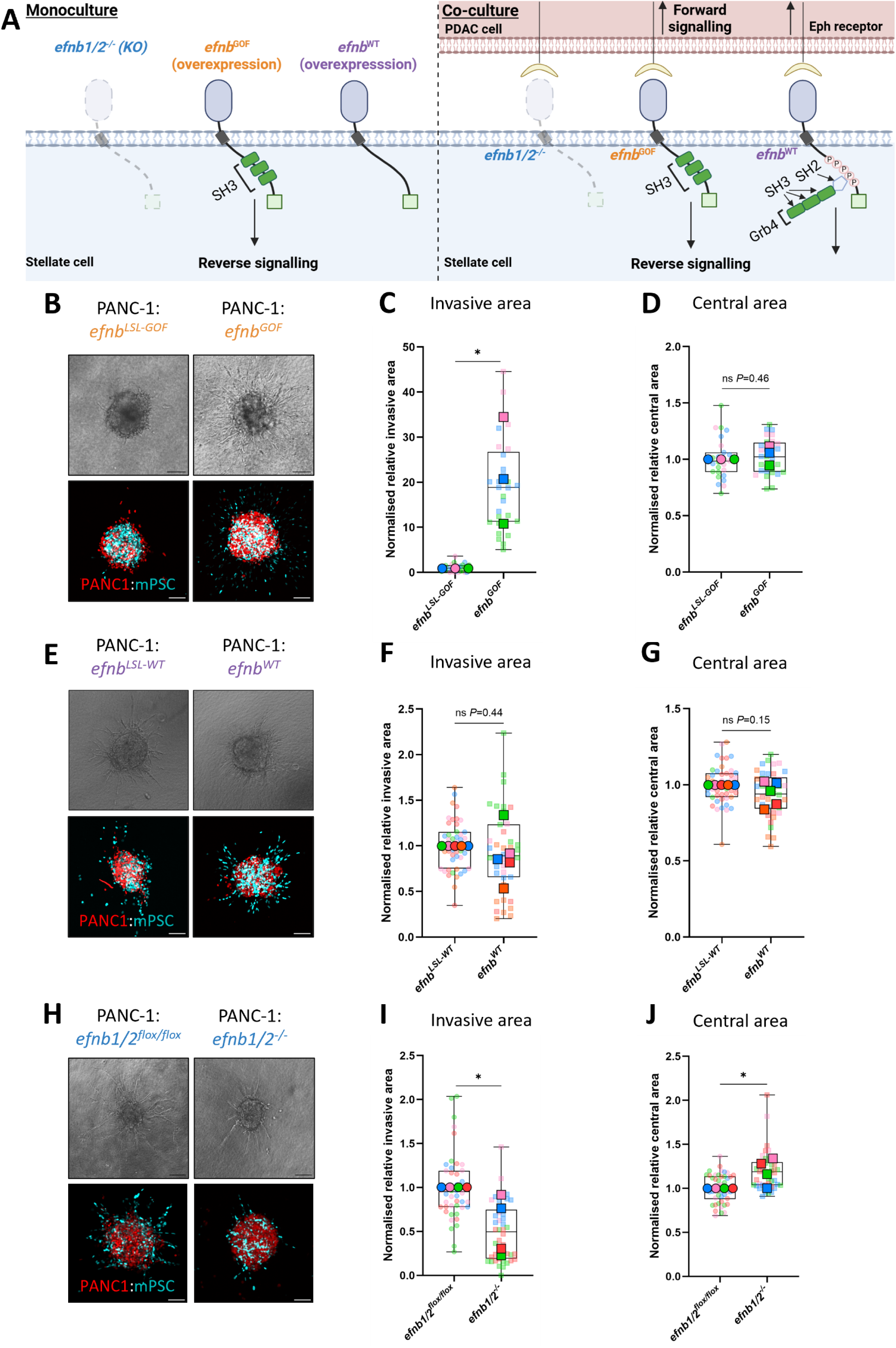
Reverse EphrinB signalling is required for CAF invasion. (A) Schematic detailing panel of genetically engineered Ephrin stellate cells, indicating effects on reverse signal. *efnb^WT^* (purple) has the normal efnb structure, whilst the *efnb^GOF^* (orange) has the backbone of efnb replaced with SH3 domains, allowing for constitutive reverse signalling. The *efnb1/2^- /-^* (blue) has a complete deletion of both *efnb1* and *efnb2.* (B, E, H) Representative brightfield and confocal spheroid images of H2B-RFP PANC-1 (red) and H2B-iRFP stellate (cyan) cells. For each condition, comparison was made to cells with the same genotype but without cre-mediated recombination. Spheroids were cultured for 2 days prior to imaging. Scale bar = 100 μm. (C, D, F, G, I, J) Spheroid quantification of invasive (C, F, I) and central (D, G, J) area normalised to the relevant control cell line. Graphs presented as superplots with 1 point = 1 spheroid, coloured by biological replicate with averages per repeat shown outlined and in bold. Unpaired two-tailed t-test (C, D, F, G) \**P*<0.05 and Mann Whitney-U test (I, J) \**P*<0.05.

For the overexpressing WT and GOF lines, as well as the *efnb1/2^-/-^* line, Cre-Lox recombination via excision of loxP flanked exons (flox) to remove the lox-stop-Lox (LSL) cassette, was required for expression of the engineered construct (**Supplementary Figure 2A**). Lines prior to Cre-recombinase expression were termed *efnb^LSL-WT^, efnb1/2^flox/flox^* and *efnb^LSL-GOF^*. Successful recombination of each line was confirmed in murine brain protein lysates following crossing to a mouse with a neural-specific Emx1-Cre driver (**Supplementary Figure 2B**). Evidence of a GOF phenotype was observed in the *efnb^GOF^*mice, with agenesis of the corpus callosum and malformation of the hippocampus, which could not be seen in the *efnb^WT^* overexpressing mouse brain (**Supplementary Figure 2C**). Successful recombination of *efnb1/2^-/-^*in the knockout was confirmed by qPCR (**Supplementary Figure 2D**).

To assess the impact of EphrinB signalling alteration on stromal-led invasion, we combined the EphrinB reverse signalling GOF stellate cells with EPHB2-expressing PANC-1 cancer cells in spheroid assays. Spheroids of *efnb^GOF^*/PANC-1 cells showed significantly increased invasion compared to spheroids containing parental stellate cells not exposed to Cre recombination (*efnb^LSL-GOF^*) (**Figure 2B-D**). In addition to constitutive reverse signalling, following recombination the GOF cells will also exhibit increased ligand capacity due to overexpression of the GOF EphrinB protein. To determine if the increased stellate cell migration was due to constitutive reverse signalling or enhanced forward signalling, we compared the action of GOF cells with stellate cells overexpressing a wild type EphrinB, driven by the same *Rosa26* promoter as the GOF construct. No difference in invasion or central area was observed between spheroids incorporating either *efnb^WT^* or parental control cells (*efnb^LSL-WT^*, **Figure 2E-G**). This confirms that the increased invasion seen for *efnb^GOF^* spheroids results from constitutively active, Grb4 SH3 domain-mediated, reverse signalling.

Consistent with a role for reverse signalling in migration, genetic deletion of EphrinB1/2 in stellate cells (*efnb1/2^-/-^*) resulted in a significant decrease in stellate/PANC-1 spheroid invasion compared to parental control (*efnb1/2^flox/flox^*) (**Figure 2H**). Interestingly, these knockout spheres also had a significantly increased central area, suggesting efnb1/2 loss either aids global cellular proliferation or, more likely, the lack of cellular invasion means more cells are present within the central spheroid (**Figure 2J**).

### EPHB2-Fc stimulation increases EPH-EphrinB2 clustering

Having shown that EphrinB2/Grb4 reverse signalling drives stromal invasion, we explored the dynamics of ligand-receptor binding. To mimic clustering of EPH-Ephrin interactions required for maximal signalling, soluble recombinant EPHB2-Fc ectodomain protein was pre-clustered with IgG^25, 27^ and then used to stimulate EphrinB2 expressing CAFs (**Figure 3A, B**). Subsequent IF labelling visualised EphrinB2/EPHB2-Fc cluster formation as distinct foci, as reported previously^25^. Specificity was confirmed through no appreciable signal detected in *efnb1/2^-/-^*cells (**Figure 3C**). Quantification of these foci allowed measurements of cluster size, cluster intensity and cluster number. Values were only counted if they were above the maximum value measured under control conditions (IgG only and serum-free medium) (**Supplementary Figure 3A-C**). To highlight the spread of our data across conditions we used superplots, which enabled statistical analysis per biological replicate, thereby avoiding oversampling^28^.

**Figure 3.**
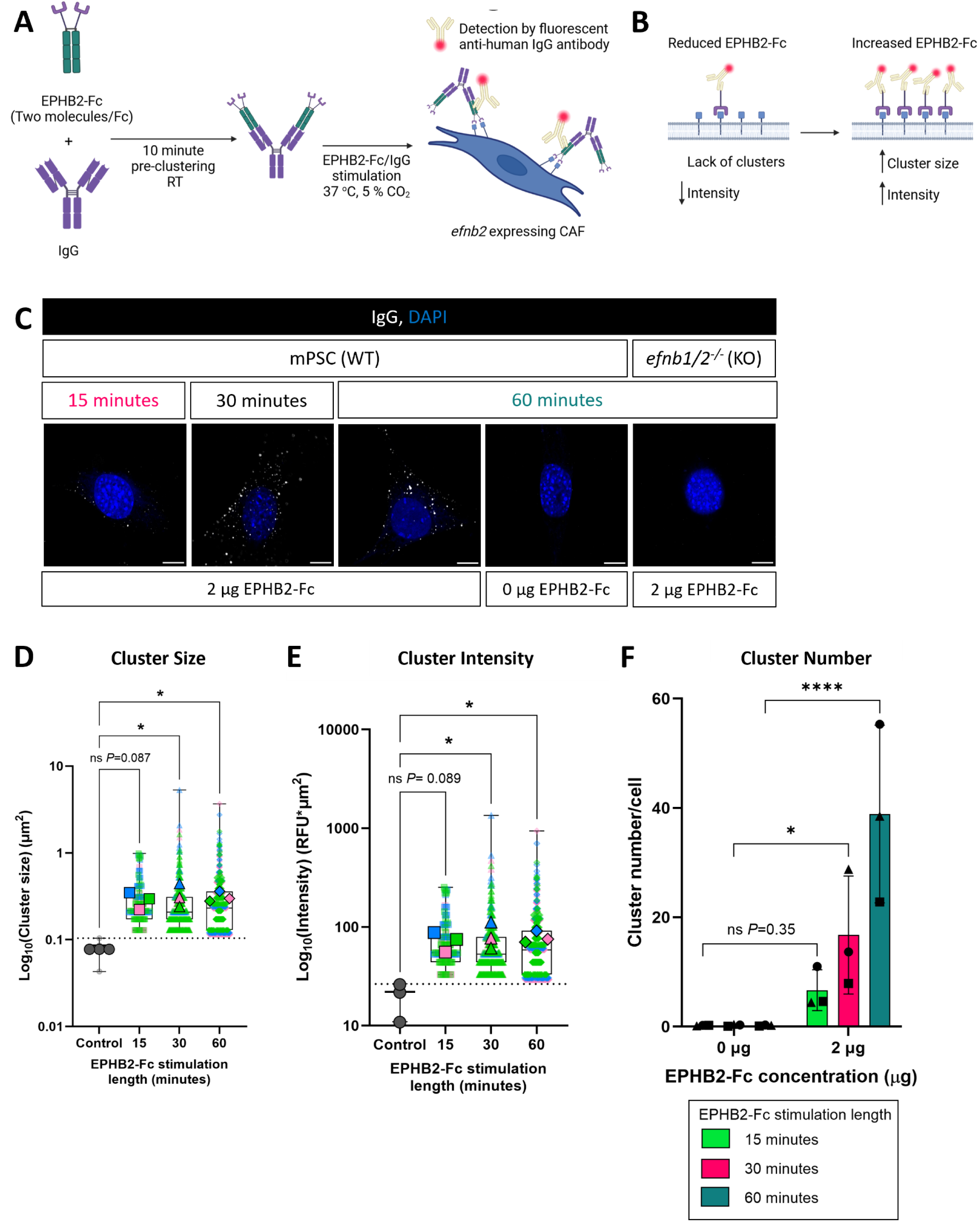
EPHB2-Fc causes time dependent increase in EPH-EphrinB clusters. (A) Schematic of CAF stimulation with recombinant EPHB2-Fc. IgG pre-clustering of EPHB2-Fc is required to ensure greater probability of successful receptor/ligand binding. Detection of ligand/receptor-Fc binding was achieved by using the anti-human IgG antibody. (B) Schematic indicating effects of cluster size on antibody labelling. Both schematics were made using Biorender.com. (C) Representative 63x confocal images of EPHB2-Fc/IgG binding (white) to EphrinB2 expressing or *efnb1/2^-/-^*CAF cells over 15, 30, and 60 minutes (blue indicates nuclei). Scale bar = 10 μm. (D-F) Cluster size (D), cluster intensity (E) and cluster number per cell (F) were quantified using the *analyse particles* macro on Fiji ImageJ. IgG alone values represented by dotted line. Graphs presented as superplots with 1 point = 1 cluster, coloured by biological replicate with averages per repeat shown outlined and in bold. Kruskal Wallis with uncorrected Dunn’s test (D, E) or two-way ANOVA with Tukey’s multiple comparisons test (F) was used \**P*<0.05, \*\**P*<0.01 (N=3).

EPHB2-Fc stimulation caused a time and concentration dependent increase in EPH-EphrinB2 cluster size, intensity and number, consistent with increasing signalling output (**Figure 3D, E, F, Supplementary Figure 3D-H**). EPHB2-Fc specificity was demonstrated through a lack of cluster formation in *efnb1/2^-/-^* stellate cells (**Figure 3C**, **Supplementary Figure 3D**). This specificity was further confirmed through western blotting for total EphrinB2, which showed no detectable expression in *efnb1/2^-/-^* stellate cell lysates (**Supplementary Figure 3I**). Thus, our data show that EPHB2-Fc can stimulate EphrinB2 on CAF cells, with improved stimulation yielded by increasing EPHB2-Fc concentration and time of stimulation.

### Pharmacological inhibition of EPH-Ephrin tetramerisation significantly reduces EPHB2-Fc/EphrinB clustering

Recent development of a tetracycline derivate, MCD, exhibiting direct EphB tyrosine kinase inhibition, has highlighted the impact of therapeutically targeting this pathway in pathologies such as chronic pain^29^. Consistent with a requirement for forward signalling in cancer invasion, MCD treatment significantly reduced CAF-led invasion in our spheroid assay (**Supplementary Figure 4**). However, our data suggest that both forward EPHB and reverse EphrinB2 signalling is important for CAF-led invasion, indicating that maximal therapeutic effect could be achieved through an agent that targets both signalling directions. To this end, we utilised a novel tetramerisation inhibitor, A20, which blocks EPHB-EphrinB binding with submicromolar IC_50_ values^21, 22^.

In an IncuCyte chemotaxis assay, A20 produced a concentration-dependent decrease in CAF (IC_50_ = 0.64 μM) and cancer cell (IC_50_ = 1.18 μM) migration towards a serum stimulus over 24 hours (**Figure 4A, B**), without affecting overall proliferation of either cell type (**Supplementary Figure 5A**). MTT assay, to assess toxicity, showed good tolerance of A20 at concentrations below 5 μM in both CAFs and cancer cells (**Supplementary Figure 5B, C**).

**Figure 4.**
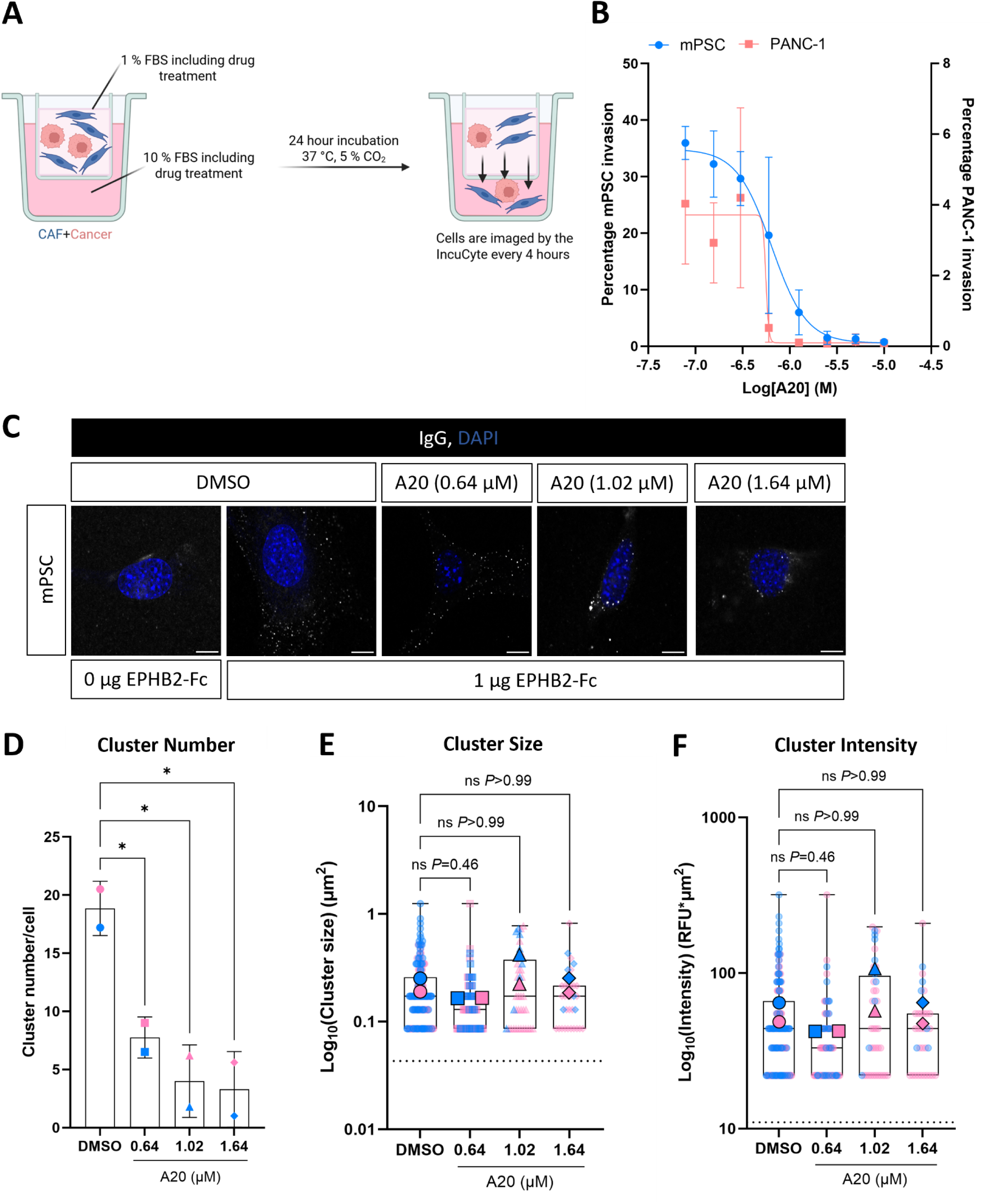
Novel EPH-Ephrin tetramerisation inhibitor significantly reduces EPHB2-Fc/EphrinB clustering. (A) Schematic for the IncuCyte chemotaxis assay. CAF and PDAC cells were plated in the apical compartment with 1 % FBS medium, with 10 % FBS added to basolateral compartment, allowing a concentration gradient. (B) Quantification of cell invasion as a percentage normalised to the initial cell number in the apical compartment for PANC-1 (red) and mPSC (blue) cells at varying concentrations of A20 (N=3). (C) Representative 63X confocal images of cluster formation following stimulation with 1 μg EPHB2-Fc for 30 minutes +/- A20 at indicated concentrations (blue indicates nuclei). Scale bar = 10 μm. (D-F) Cluster number per cell (D), cluster size (E) and cluster intensity (F) were quantified using the *analyse particles* macro on Fiji ImageJ. IgG alone values represented by dotted line. Superplots show all data above threshold (dotted line), with 1 point = 1 cluster, coloured by biological replicate with averages per repeat shown outlined and in bold. Ordinary one-way ANOVA with Dunnett’s multiple comparisons test (D) and Kruskal Wallis with Dunn’s multiple comparisons test (E, F) utilised \**P*<0.05 (N=2).

We next examined whether A20 could block EPHB2-Fc/EphrinB2 binding, reasoning that if A20 blocked tetramer formation, we should observe a significant decrease in cluster size, intensity and number of clusters. EphrinB2-expressing CAFs were stimulated with 1 μg of pre-clustered EPHB2-Fc for 30 minutes, together with either DMSO or A20. Concentrations of A20 were chosen from the IncuCyte chemotaxis assay that represented IC_25_ (0.64 μM), IC_50_ (1.02 μM) and IC_90_ (1.64 μM) values for inhibition of invasion. A20 produced a concentration-dependent decrease in cluster number, with a significant decrease in EPH-EphrinB2 cluster formation down to 0.64 μM (**Figure 4C, D**). Interestingly, when analysing the size and intensity of any remaining clusters, no change could be seen compared to DMSO (**Figure 4E, F**). This suggests A20 can block EPHB2-Fc/EphrinB2 binding.

As A20 can block bidirectional signalling, we investigated whether A20 blocks forward signalling in EPHB-expressing cancer cells stimulated with EPHRINB2-Fc. As with CAFs, EPHRINB-Fc stimulation of EPHB2-expressing cancer cells promoted cluster formation (**Supplementary Figure 5D**). Accordingly, A20 caused a significant reduction in cluster number but not cluster size or intensity (**Supplementary Figure 5E-G**).

### Small molecule inhibition of Ephrin signalling significantly decreases PDAC invasion

Having confirmed that A20 can reduce the ability of EPHB2-Fc to bind and cluster EphrinB2 on CAFs, we assessed whether A20 could affect CAF-led invasion using our spheroid model. Initial experiments using MIA PaCa-2 PDAC cells and mouse CAFs, showed that A20 administration caused a significant decrease in invasion, with no discernible toxicity (**Figure 5A-C**).

**Figure 5.**
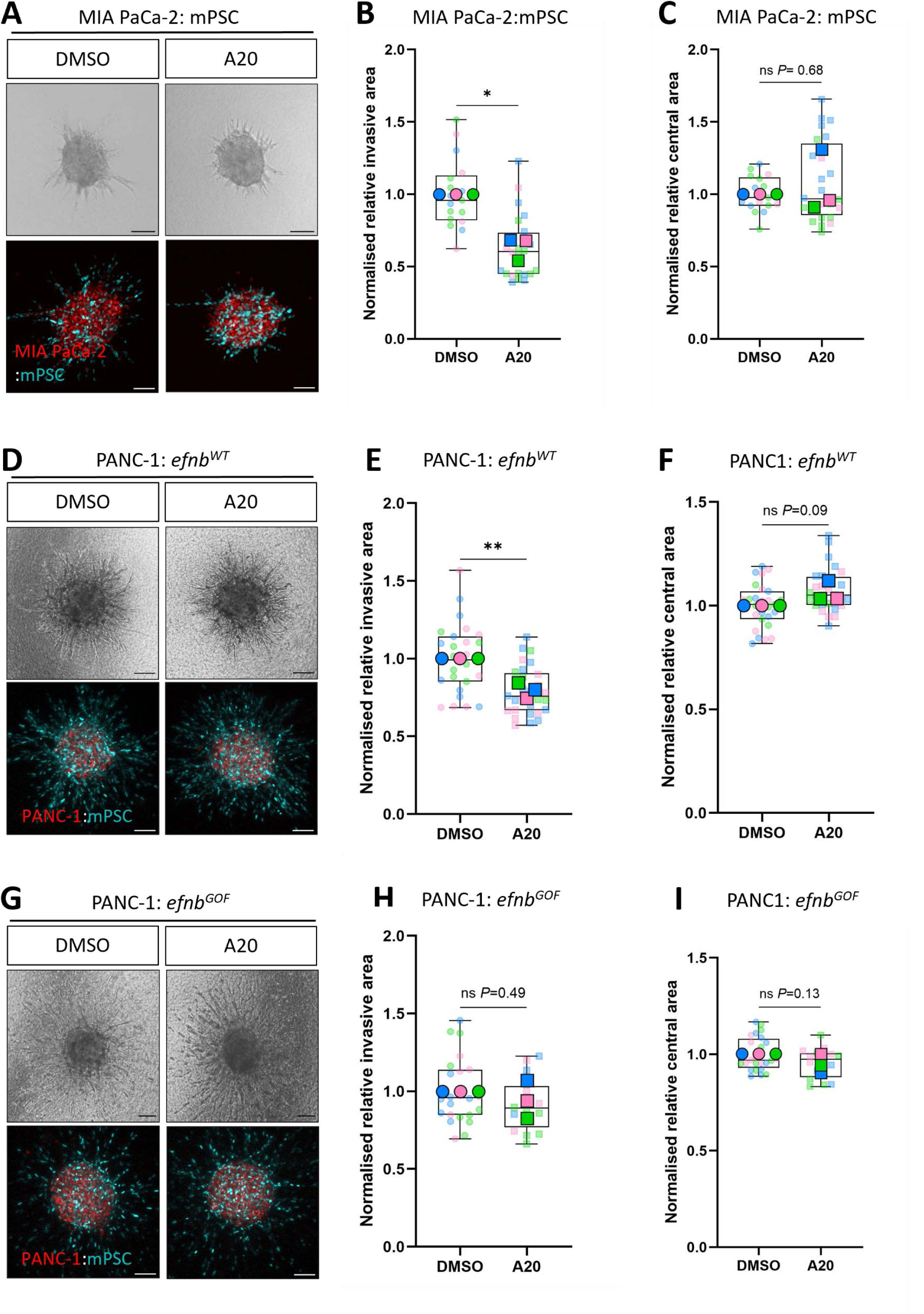
Small molecule inhibition of Ephrin signalling significantly decreases PDAC invasion. (A) Representative brightfield and confocal spheroid images of H2B-RFP MIA PaCa-2 (red) and H2B-GFP CAF (cyan) cells. Spheroids were cultured for 3 days prior to imaging, with replenishment of medium containing DMSO or A20 (10 μM) every 24 hours. Scale bar = 100 μM (N=3). (B, C) Spheroid quantification of invasive (B) and central (C) area comparing DMSO to A20 (10 μM). Unpaired two-tailed t-test utilised \**P*<0.05 (N=3). (D, G) Representative brightfield and confocal spheroid images for H2B-RFP PANC-1 (red) and H2B-iRFP stellate (cyan) cells (either *efnb^WT^* (D) or *efnb^GOF^* (G)). Spheroids were cultured for 2 days with daily replenishment of medium containing DMSO or A20 (3 μM). (E, F, H, I) Spheroid quantification of invasive (E, H) and central (F, I) areas normalised to the relevant control cell line. Superplots are shown with 1 point = 1 spheroid, coloured by biological replicate and averages per repeat shown outlined and in bold. Unpaired two-tailed t-test utilised \*\**P*<0.01 (N=3).

To understand whether EPHB2 expression level affected response to A20, we identified a high EPHB2 expressing PDAC cell line (PANC-1, **Supplementary Figure 6**) to co-culture with *efnb^WT^*stellate cells. The *efnb^WT^* stellate cells overexpress EphrinB, so we reasoned it would be optimal to assess A20 efficacy with a high expressing EPHB2 cell line and an overexpressing EphrinB cell line, where activation of both forward and reverse signalling will be increased. A20 treatment decreased global spheroid invasion significantly, with both cell types affected despite increased ligand capacity of *efnb^WT^* cells, highlighting the ability of A20 to block EPHB/EphrinB interactions (**Figure 5D-F**).

We have shown that induced constitutive reverse Grb4 SH3 domain-mediated signalling of *efnb^GOF^* stellate cells caused a significant increase in spheroid invasion (**Figure 2B-D**). To confirm that A20 was acting specifically to block EPHB2/EphrinB signalling, we treated *efnb^GOF^*/PANC-1 spheroids, which we hypothesised would be unresponsive to A20, as reverse signalling is constitutive and ligand-independent in these cells. As predicted, *efnb^GOF^*/PANC-1 spheroids showed no significant change in invasive or central area after A20 treatment (**Figure 5G-I**).

### Ephrin inhibition blocks local PDAC invasion *in vivo*

Having confirmed that A20 blocked EPHB2/EphrinB binding, significantly decreased spheroid invasion, and exhibits good pharmacokinetic properties^21, 22^, we examined the effects in mouse models of PDAC. The mouse PDAC cell line TB32048, derived from the *Kras^G12D/+^*; *Trp53^R172H/+^*; *Pdx1-Cre* (KPC) mouse model^30^, was orthotopically implanted into recipient C57BL/6 mice pancreata (**Figure 6A**). After 7 days, the mice were injected intraperitoneally (IP) with either PBS or A20 (10 mg/kg), three times per week. At day 24, the pancreatic tumours were measured, dissected, sectioned and stained with Haematoxylin and Eosin (H&E). Primary tumour mass and volume showed no significant differences between PBS and A20 treated mice (**Figure 6B, C**). This corroborated *in vitro* observations, where A20 concentrations below 5 μM were able to block invasion without causing growth arrest or cell death. Interestingly, H&E staining of pancreata revealed A20 treated mice exhibited an absence of cancer cell invasion into the adjacent, normal pancreatic tissue compared to the vehicle dosed mice (**Figure 6D**). It was apparent that A20 treatment led to a compartmentalisation of the tumour, with a distinct boundary between normal pancreatic acini and tumour tissue, in comparison to the vehicle controls, where normal pancreatic acini were clearly overwhelmed by invading tumour tissue (**Figure 6D**). Quantification of local invasion highlighted a significant decrease in A20-treated mice compared to controls (**Figure 6E**).

**Figure 6.**
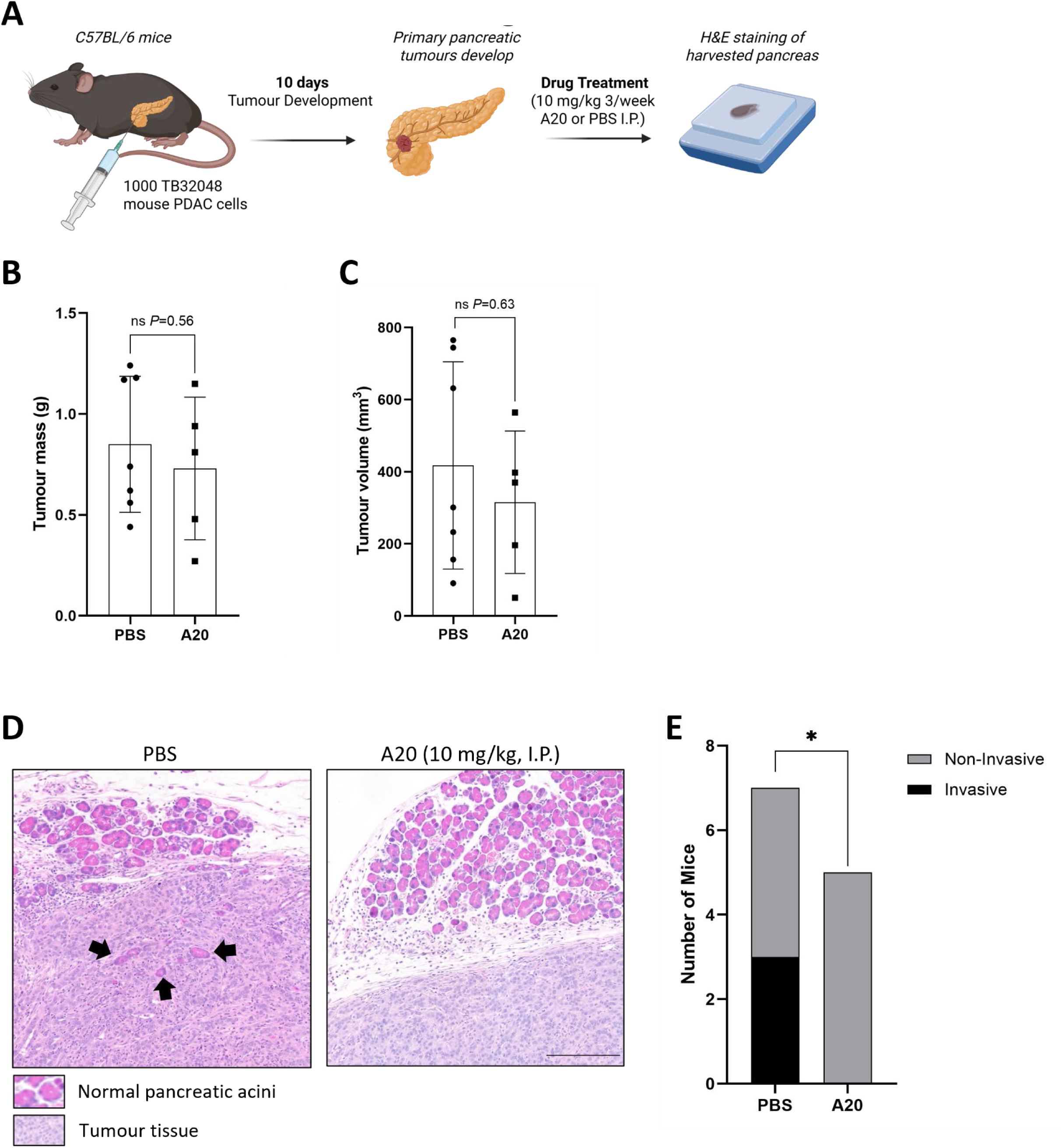
Ephrin inhibition blocks local PDAC invasion *in vivo*. (A) Schematic overview of primary disease model. TB32048 PDAC cells were orthotopically injected into C57BL/6 mice. After 10 days of tumour development, I.P. injection of PBS or A20 was given (10 mg/kg x3/week) until day 24. (B, C) Quantification of primary tumour mass and volume respectively per mouse. Statistical analysis by either two-tailed unpaired t-test (B) or Mann-Whitney U test (C), ns denotes *P*>0.05 (N=7 PBS and N=5 A-20). (D) H&E staining of pancreas sections taken from PBS control and A20 treated mice. Arrows indicate local invasion in pancreas from vehicle treated animals. (E) Quantification of local invasion per mouse taken from histological sections. Statistical analysis by unpaired two-tailed, t-test, \**P*<0.05 (N=7 PBS and N=5 A-20).

### Ephrin inhibition blocks metastasis

As the majority of PDAC patients present to the clinic with metastatic disease^31^, we next examined the effect of A20 in an *in vivo* model of advanced disease. For this work, we utilised the integrin β6^+^ TB32048 PDAC cell line, which has greater metastatic potential *in vivo* than the parental TB32048 cancer line^32^. Transcriptomic analysis of β6^+^ TB32048 cells revealed an increase in expression of Ephb2 and a significant downregulation of EphrinB2 compared to parental TB32048 cancer cells (**Supplementary Figure 7A**).

Syngeneic C57BL/6 mice were injected orthotopically with 1000 β6^+^ TB32048 PDAC cells and housed for 7 days to allow tumour development (**Figure 7A**). Tumour growth was monitored by IVIS imaging, measuring total and fold change flux (**Supplementary Figure 7B-D**). Mice were injected IP with either PBS or A20 (10 mg/kg) three times per week for 4 weeks.

**Figure 7.**
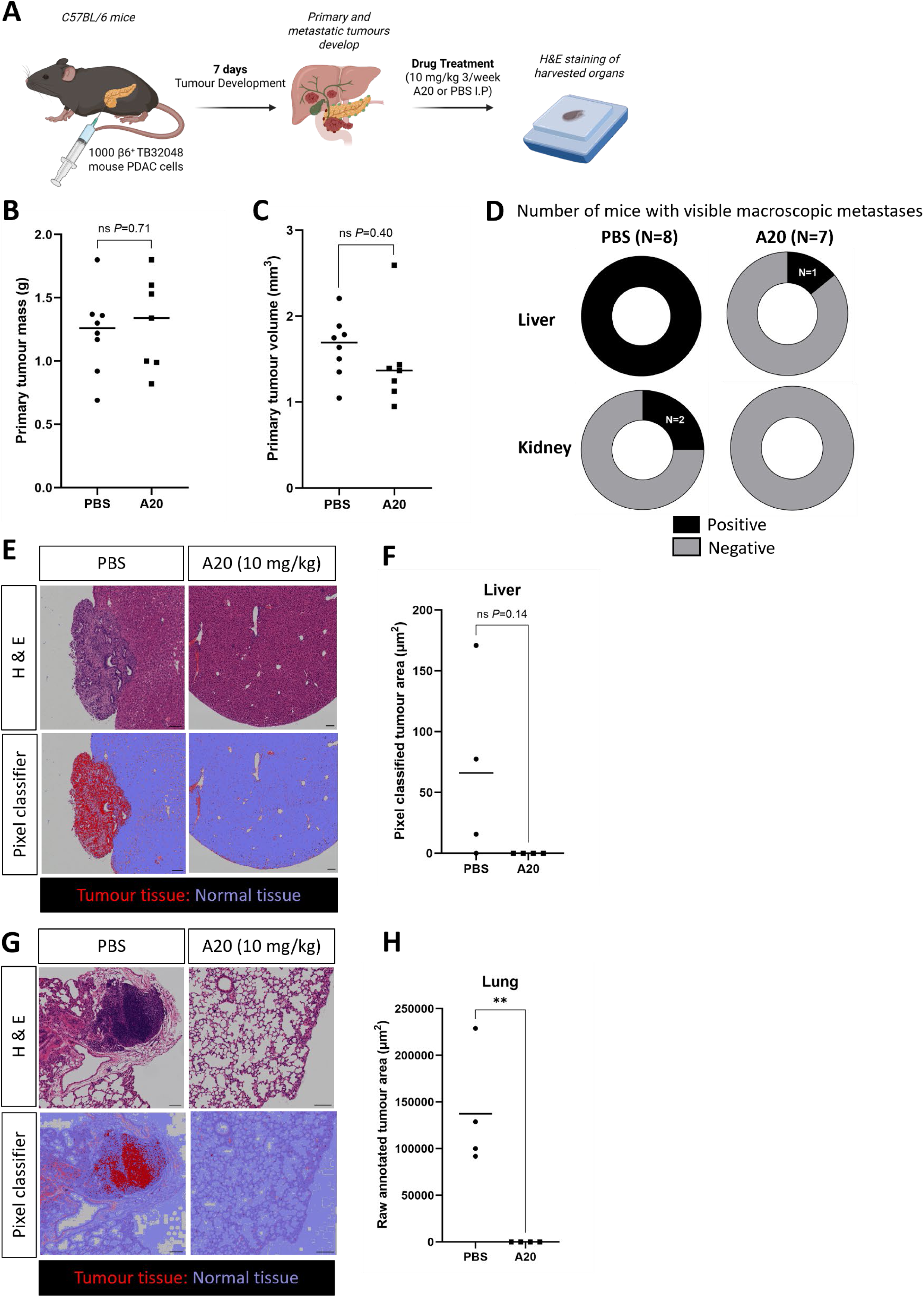
Ephrin inhibition blocks metastatic tumour formation. (A) Schematic overview of experimental model. β6^+^ TB32048 PDAC cells were injected orthotopically into isogenic C57BL/6 mice. Tumours developed in 7 days as confirmed by IVIS imaging, after which IP injection of A20 was given (10 mg/kg x3/week) for 4 weeks. (B, C) Quantification of tumour mass and volume respectively per mouse for primary tumours. Statistical analysis by two-tailed unpaired t-test, ns denotes *P*>0.05 (N=8 PBS, N=7 A20). (D) Number of visible macroscopic metastatic tumours in organs harvested from both PBS and A20 treated mice (N=8 PBS or N=7 A20). (E) H&E staining of liver sections from PBS and A20 treated mice (N=4/group), with QuPath pixel classifier trained below to recognise tumour tissue (red) vs. normal hepatocytes (blue). (F) Quantification from pixel classification of liver tissues. Two-tailed unpaired t-test ns *P*>0.05 (N=4/group). (G) H&E staining of lung tissue taken from PBS or A20 treated mice (N=4/group), with QuPath pixel classification below showing an inability of the software to recognise tumour (red) vs. normal tissue (blue). (H) Quantification of raw tumour area annotated in QuPath. Two-tailed unpaired t-test \*\**P*<0.01 (N=4/group).

Consistent with our previous experiment, primary tumour mass and volume were not appreciably different between treatment conditions (**Figure 7B, C**). However, there was a marked decrease in the number of organs with visible macroscopic metastases in A20-treated mice (**Figure 7D**). Tumour formation in spleen, peritoneum and intestine was discounted as metastases, due to proximity to the pancreas suggesting local spread at the time of orthotopic injection. However, tumour growth at these sites was reduced with A20 treatment (**Supplementary Figure 7E**). In contrast, the “true” metastases in liver and lung showed no discernible metastases in both organs at dissection for A20 treated mice (**Figure 7E-H**), highlighting the potential of A20 to block disease progression.

## Discussion

Deepening our knowledge of PDAC progression is of paramount importance to develop more efficacious, targeted treatments. PDAC is characterised by a desmoplastic stroma, making treatment delivery challenging^2^. CAFs dominate the PDAC microenvironment and are widely regarded as a driving force in ECM production^33, 34^, treatment resistance^33, 35^ and PDAC invasion ^33^. Here, we have shown that the EPHB2-EphrinB2 signalling axis can mediate PDAC-CAF crosstalk during invasion, highlighting this pathway as a therapeutic target.

Public datasets, based on transcriptomic analysis of bulk pancreatic cancer patient samples, indicated that upregulation of *EFNB2* correlated with worse overall survival. Positive correlation of EPHRINB2 expression with worsened overall survival has been observed in a range of cancers including PDAC^36^, head and neck squamous cell carcinoma^36^ and bladder urothelial carcinoma^36, 37^, underscoring the potential for EPHRINB2 as a biomarker and therapeutic target. Upregulation of EPHRINB2 in pancreatic cancer has been previously reported utilising the same datasets^11^. However, the authors proceeded to validate EPHRINB2 in PDAC cell lines which express lower levels of EPHRINB2 mRNA or protein relative to stromal components such as CAFs -which would be present within bulk tumour analysis^11^. Here, we identified EphrinB2 as a CAF-expressed molecule important for cancer cell invasion.

Our results indicate a driving role of CAF-EphrinB2 in directing PDAC cell migration and invasion (**Figure 8**). Using stellate cells harbouring a GOF EphrinB protein, we have further shown that constitutive EphrinB reverse signalling in stellate cells leads to a significant increase in spheroid invasion. We also observed a dependence of cancer cell EPHB2 for CAF-led invasion, mirroring similar observations in prostate cancer invasion^15^. Together, this indicates that both forward signalling through cancer cells, and reverse signalling through CAFs is required for invasion^38, 39^. Targeting both forward and reverse signalling would therefore be needed for maximal therapeutic effect. Hence, we explored the use of newly developed EPH-Ephrin tetramerisation inhibitors that block both forward and reverse signalling. The tetramer inhibitor A20 successfully blocked 3D spheroid invasion, decreasing CAF-led cancer cell invasion with minimal effect on cellular proliferation at concentrations lower than 5 µM. Extending these observations *in vivo* we observed a block on local invasion in primary disease, and a block on metastatic tumour formation in advanced disease (**Figure 8**). This work presents a promising new approach for PDAC therapy that could be beneficial at the primary and metastatic tumour level.

**Figure 8.**
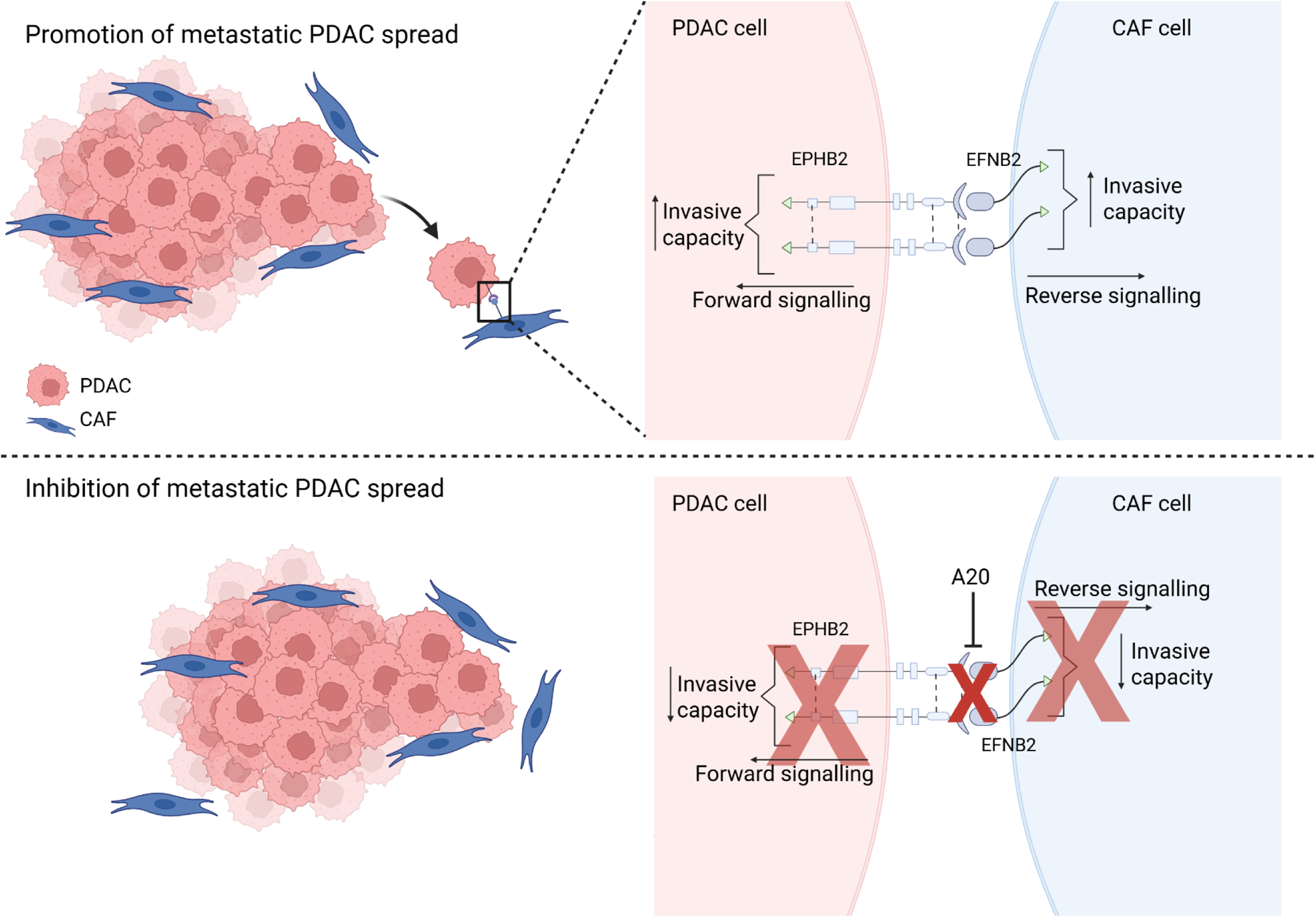
Eph/Ephrin signalling promotes CAF-led PDAC invasion and metastasis. Schematic made using Biorender.com.

This work is the first of its kind allowing efficacious blockade of forward and reverse signalling. Initial attempts to target this signalling family have utilised monoclonal antibodies (mAbs), raised against either EPHRIN ligands or EPH receptors. Many mAbs developed have focused on binding to EPH receptors. For example, 1C1 was developed as a human anti-EPHA2 mAb^40^, with an antibody-drug-conjugate of 1C1 (MEDI-547) advancing to clinical trials for solid tumours^41^. Although binding on target, this conjugate compromised safety after administration, causing internal bleeding and coagulation, leading to early termination of the Phase 1 trial^41^. Our work, and other *in vivo* studies by our colleagues^21, 22^, have indicated Eph/Ephrin tetramerisation inhibitors are well tolerated.

In conclusion, we have shown that EPHRINB2 reverse signalling can drive CAF invasion, which promotes cancer cell invasion through cancer cell EPHB2. Disruption of this bidirectional receptor-ligand signalling reduces invasion both *in vitro* and *in vivo*, critically at the primary and advance disease setting. These data present EPHB2-EPHRINB2 inhibition as a novel therapeutic strategy for PDAC patients and supports the continued clinical development of tetramerisation inhibitors.

## Methods

### Generation of Cre-inducible mice overexpressing efnb^WT^ or efnb^GOF^

Mice were engineered to constitutively overexpress either a Flag epitope-tagged WT EphrinB protein or a reverse signalling gain-of-function (GOF) protein, conditional upon Cre-mediated recombination. To accomplish this, CRISPR was used to directly target WT and GOF constructs into the *Rosa26* locus in mouse embryos and mice were recovered that transmitted the desired insertions through the germ line.

Specifically, a single guide RNA (sgRNA) molecule (sgRosa26-1) was used to generate double strand breaks in the *Rosa26* locus. Embryos were injected with sgRosa26-1, Cas-9 mRNA/protein, and targeting vectors for recombination into the *Rosa26* locus, and then transferred into pseudopregnant females to allow full development and birth of pups. Targeting vectors containing 5’ and 3’ arms of homology to the *Rosa26* locus that flank the region targeted by sgRosa26-1 were co-injected with the sgRNA/Cas-9 mixture to mediate homology-directed repair of the *Rosa26* locus. This led to insertion of sequences containing a CAG promotor-loxP-STOP-loxP expression cassette containing an open reading frame to express a Flag-tagged version of either the efnb^WT^ protein or the efnb^GOF^ constitutive-active protein, after exposure to Cre recombinase. Inclusion of an Internal Ribosome Entry Site (IRES) facilitated co-expression of green fluorescent protein (GFP) with the Flag-tagged EphrinB proteins.

To confirm GOF activity, the WT and GOF lines were crossed to mice containing an *Emx1-Cre* driver to delete the loxP-STOP-loxP element and induce expression of the engineered EphrinB proteins and an IRES-linked GFP in radial glia and excitatory neurons in the neocortex.

### Cell culture

All cell lines were cultured in Dulbecco’s Modified Eagle Medium/Nutrient Mixture F-12 (DMEM F-12; 11330-032 Gibco Thermo Fisher Scientific) supplemented with 10 % Foetal Bovine Serum (FBS; A5256801 Gibco Thermo Fisher Scientific) and 1 % penicillin/streptomycin (15140-122 Gibco Thermo Fisher Scientific). Cells were maintained at 37 °C in a 5 % carbon dioxide (CO_2_) humidified incubator and regularly tested for mycoplasma.

Pancreatic stellate cells harbouring *efnb^LSL-WT^*, *efnb^LSL-GOF^* and *efnb1/2^flox/flox^* genetic constructs were isolated as previously described^42^, with isolated cells immortalised by transduction with pBabe SV40 large T plasmid^24^. For all three cell lines, Cre-mediated excision was required to generate specific Ephrin mutant phenotypes (efnb^WT^, efnb^GOF^, efnb1/2^-/-^). This was achieved through 24-hour adenoviral transduction of active Cre (6 x10^10^ PFU p/mL; kindly provided by Dr Rita Pedrosa, (Queen Mary University of London; QMUL).

The MIA PaCa-2 and PANC-1 PDAC lines, and immortalised human pancreatic stellate cells (PS1) were provided by Professor Hemant Kocher (QMUL). Immortalised mouse pancreatic stellate cells (mPSC) were isolated as described previously ^24^. Fluorescent constructs utilised for PDAC, CAF and stellate cell lines, as indicated in figure legends, were provided by Professor Susana Godinho (QMUL).

The parental TB32048 and integrin β6^+^ TB32048 pancreatic cancer cell lines are through an *FSF-Kras^G12D/+^*; *LSL-Trp53^frt/+^*; *Pdx-1-Flp* genetic background, with a construct for the overexpression of GFP and luciferase by the pHAGE PGK-GFP- IRES-LUC-W plasmid, developed by Professor Darrell Kotton (Boston University, USA) ^43^. The development of these cell lines and corresponding mouse models has been previously described^32^.

### Real-time quantitative polymerase chain reaction

The Monarch Total RNA miniprep kit (#T2010S, New England BioLabs) was used to extract RNA, following manufacturer’s instructions, with RNA quantified using a NanoDrop^TM^ One/One^C^ Microvolume UV-Vis Spectrophotometer (Thermo Fisher Scientific). Reverse transcription (RT) was performed using LunaScript RT SuperMix Kit (M3010L, New England BioLabs) following manufacturer’s instructions. The resulting complementary DNA (cDNA) was amplified through quantitative polymerase chain reaction (qPCR) using Luna Universal qPCR Master Mix (M3033G; New England BioLabs) and relevant primers (**Supplementary Table 1**). The StepOne Plus real-time PCR machine (Thermo Fisher Scientific) was used following manufacturer’s instructions. Relative gene expression was calculated using the 2^-ΔΔCT^method^44^, normalising to β-actin control. To confirm PCR product sizes, and lack of contamination in negative control reactions, samples were run on a 2 % (w/v) agarose gel containing GelRed (41003, Biotium), imaged using a UV trans-illuminator (UVP97063001, Thermo Fisher Scientific).

### EPH/Ephrin stimulation

Recombinant EPHB2-Fc protein (5189-B2) and recombinant EFNB2-Fc (496-EB) were purchased from R&D systems.

After serum starvation, recombinant Fc protein (200 μg/mL) was pre-clustered for 10 minutes at room temperature (RT) by incubation with goat human-Fc protein (1200 μg/mL; 31470, Thermo Fisher Scientific). Stimulation was for 0, 15, 30 or 60 minutes at 37 °C and 5 % CO_2_, after which the stimulant was removed, and the cells were processed for downstream applications.

### Western blotting

Brains from EphrinB genetically modified mice (*efnb^LSL-WT^*, *efnb^WT^*, *efnb^LSL-GOF^* and *efnb^GOF^*) were lysed with buffer containing 50 mM Tris-HCl, pH 7.5, 200 mM NaCl, 5 mM MgCl_2_, 1 % NP-40, 10 % Glycerol, 1 mM Dithiothreitol (DTT), 1 mM Phenylmethylsulfonyl fluoride and protease inhibitor mixture (Roche Molecular Biochemicals). Stellate and CAF cells were lysed in Pierce radioimmunoprecipitation assay (RIPA) lysis and extraction buffer (89900, Thermo Fisher Scientific), with added 1 mM protease inhibitor cocktail (P8465, Sigma-Aldrich), 10 mM sodium fluoride (Sigma-Aldrich) and 1 mM sodium orthovanadate (Sigma-Aldrich).

Protein denaturation was performed at 65 ֯C for 5 minutes for mouse brain lysates, or 95 ֯C for 5 minutes for stellate and CAF cell lysates. Protein samples (20 μg) were separated on a 12 % SDS-PAGE gel, followed by transfer to nitrocellulose membrane (GE10600014, Sigma-Aldrich) and blocking for 45 minutes in 2 % bovine serum albumin, diluted in Tris-buffered saline with 0.1 % tween-20 (BSA-TBST; 20-7301-10, Severn Biotech and 6636848, VWR Chemicals, respectively). Subsequently, the membrane was incubated overnight at 4 °C with primary antibody including the anti-FLAG and anti-GFP antibodies (both purchased from Sigma), total EphrinB2 (APREST71886, Atlas Antibodies, Sigma-Aldrich) antibody or HSC70 housekeeping antibody (sc7298, Santa Cruz Biotechnologies); all were diluted in 5 % BSA/TBST. The next day, one hour incubation with species-appropriate horseradish peroxidase (HRP)-conjugated secondary antibody (P0448/P0447, Agilent), diluted in 5 % BSA/TBST, was performed. Immobilon® Western Chemiluminescent HRP Substrate (WBKLS0050, Sigma-Aldrich) was warmed to RT and added to the membrane for 5 minutes, prior to visualisation using the ChemiDoc^TM^ imager (Bio-Rad).

### Immunofluorescence staining

Cells grown on coverslips were fixed using 4 % paraformaldehyde/PBS (PFA; 28908, Thermo Fisher Scientific) for 20 minutes at RT. Blocking was completed with 5 % BSA/phosphate buffered saline (PBS) for 30 minutes at RT. The Alexa Fluor 647 anti-human antibody (A-21445, Thermo Fisher Scientic) was diluted in 5 % BSA/PBS and incubated for 1 hour at RT, after which the coverslips were washed and placed onto slides with ProLong^TM^ Gold Antifade Mountant with DAPI (P36931, Thermo Fisher Scientific) and allowed to dry at RT overnight.

Imaging was performed using an LSM880 or LSM710 Zeiss confocal microscope with 40X and 63X oil immersion objective.

### Immunofluorescence image quantification

Imaging files were analysed using Fiji ImageJ^45^. After setting the threshold per image to enable any clear signal to be included, *analyse particles* was set with pixel size maintained at 0-infinity, circularity set at 0.00-1.00 and outlines included, allowing measurement of integrated density and size for individual clusters through the region of interest (ROI) manager.

### Hanging drop spheroid model

Matrix-embedded heterocellular cancer/CAF spheroids where prepared as previously described^7, 24^. Briefly, cancer/CAF spheroids (1:2 ratio) were cultured with 1.2 % methylcellulose (M0512-100G, Sigma-Aldrich) for 24 hours prior to embedding in an organotypic gel, containing 2 mg/mL rat tail collagen (354249, Corning) and 17.5 % Matrigel basement membrane matrix (354234, Corning). The embedded spheroids were left for 2-3 days to allow invasion to occur. For drug treated spheres, the medium was changed daily.

Spheroids were imaged using a Zeiss Axiovert 135 (20 X magnification) before fixation in 10 % formalin (BAF-0010-25A, CellPath). Fixed spheroids were placed onto slides using mowiol (Sigma-Aldrich), and Z-stack images were taken using either the LSM710, LSM880 or LSM800 Zeiss confocal microscope (20 X objective).

Relative invasion and central sphere area were calculated using Fiji/ImageJ^46^. The following equation was used to calculate percentage invasion which was normalised to applicable control measurements per experiment:

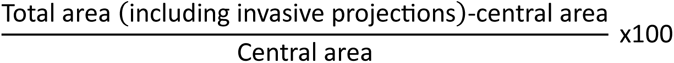

### IncuCyte chemotaxis assay

To measure cellular chemotaxis, IncuCyte® Clearview 96-well Microplate for Chemotaxis (4582, Sartorius) were imaged in an IncuCyte® S3 Live-Cell Analysis System (4647, Sartorius) every 4 hours for 24 hours. Analysis was carried out following manufacturer’s instructions utilising the IncuCyte® chemotaxis software.

### Viability assay

Cell viability was measured by 3-(4, 5-dimethylthiazolyl-2)-2, 5-diphenyltetrazolium bromide (MTT) assay. Cells were assessed at 72 hours using manufacturer’s guidelines for MTT addition (M5655-1G, Sigma-Aldrich). Absorbance was measured at 570 nm using the FLUOstar Omega filter-based multi-mode microplate reader (BMG Labtech).

### Animal experiments

All experiments were performed under UK Home Office guidelines at the Biological Services Unit (QMUL Charterhouse Square, license number PP4475211).

C57BL/6 mice were purchased from Charles River, with orthotopic injection of PDAC cells completed into the pancreas of each mouse following acclimatisation. Tumour formation was monitored by IVIS imaging once per week for the duration of the experiment. After 7-10 days, all mice were started upon IP injections of either PBS or reconstituted A20 (2.5 mg/mL resuspended in PBS to enable a 10 mg/kg dose). IP injections were carried out three times per week for 24 days or 4 weeks. Mice were checked daily to ensure no behavioural changes and weighed twice per week, with primary tumours not allowed to exceed 1.44 cm^3^. Mice were euthanised by cervical dislocation and any tumours present were collected, prepared and sectioned for H&E staining (by Pathology service; QMUL). H&E slides were scanned using a NanoZoomer S210 slide scanner on brightfield, and the raw files were uploaded to QuPath. Tumour classification was enabled by training the pixel classifier to recognise the difference between tumour and normal tissue for all organ slides with help from Dr Mike Allen (QMUL).

### Bioinformatic analysis

Data from The Cancer Genome Atlas (TCGA) pancreatic cancer cohort (PAAD; N=179) and Genotype Tissue Expression project (GTEx) (N= 171) were taken from Genome Expression Profiling and Interactive Analysis (GEPIA). Normal TCGA and GTEx data were matched, with the data enabling plotting of *EFNB2* and *EPHB2* expression in tumour bulk and normal tissue samples. The preset GEPIA parameters were kept, which included a Log_2_ fold change (Log_2_FC) cutoff of 1 and a *P*-value cutoff of 0.01.

Patient overall survival statistics including hazard ratio and log rank *P*-values for each Eph and Ephrin were acquired from Kaplan-Meier plotter web tool (Km plotter) ^47^. RNA-seq data from 177 pancreatic cancer patients were used with analysis parameters kept constant and the auto select best cut off parameter utilised.

### Statistical Analysis

GraphPad Prism version 10.6.0 was used with relevant statistical analyses indicated in figure legends and error bars indicating ± SD. For superplots, each individual point was plotted either representing a spheroid or receptor/ligand cluster (indicated in figure legends), with the colours indicating biological replicates. The average of each biological replicate is represented by outlined, bold points, with these averages utilised for any statistical tests performed. Normalisation tests were conducted to determine the use of parametric or non-parametric tests where appropriate. Unless otherwise stated, *P* < 0.05 was considered significant and denoted by one asterisk (*), whilst ** denotes *P* < 0.01, *** denotes *P* <0.001 and **** denotes *P* < 0.0001.

## Acknowledgements

We thank the Microscopy and Pathology Core Facilities at Barts Cancer Institute for expert advice and staining. The core facilities are supported by a CRUK City of London Centre Award (CTRQQR-2021\100004). We thank Professor Hemant Kocher for providing pancreatic cancer cell lines and human PS1 cells, and Professor Susana Godinho for providing fluorescent histone constructs. CLB was funded by the London Interdisciplinary Doctoral (LIDo) Programme funded by the Biotechnology and Biological Sciences Research Council (BB/T008709/1). This work was supported in part by grant MR/W006308/1 for the GW4 BIOMED2 DTP, awarded to the Universities of Bath, Bristol, Cardiff and Exeter from the Medical Research Council (MRC)/UKRI. The work by FS, HW and MH was funded by DOD/U.S. Army Medical Research and Development Command awards (W81XWH-14-1-0220, W81XWH-21-1-0949). This work was also funded by a Cancer Research UK Centre Grant to Barts Cancer Institute (C355/A25137).

## Author contributions

CLB, EPC and RPG conceived and designed the study with input from all authors. EPC performed chimeric heterocellular spheroid RNA-sequencing and siRNA spheroid experiments, together with RO’S. CLB performed patient data and cell line analysis, spheroid experiments relating to Ephrin mutant constructs and A20 optimisation/administration, CAF immunofluorescence and all antibody experiments with help from AT and EPC. FS developed Cre-inducible Ephrin WT and GOF overexpressing mice, with EPC isolating pancreatic stellate cells and CB preparing the cells for culture, which included Cre transduction. HW performed chemical synthesis for A20, with MH leading mouse models and A20 development. *In vivo* work presented for A20 was completed by EPC, SM, AJMC, GF-M, CLB, JFM and RPG. CLB, EPC and RPG wrote the manuscript with review and approval from all authors.

---

**Supplementary Figure 1.**
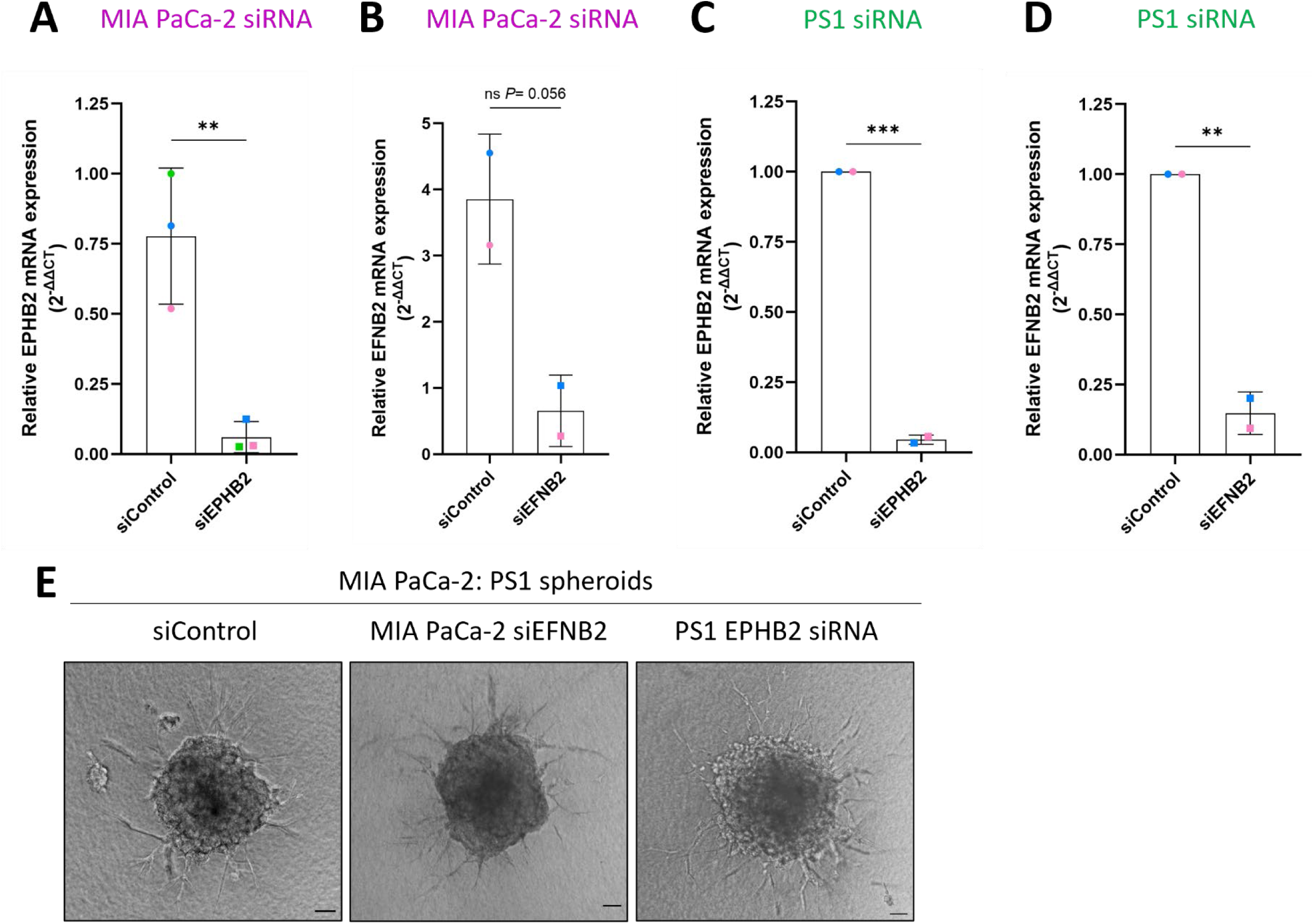
(A-D) qPCR quantification of EPHB2 and EPHRINB2 expression in MIA PaCa-2 and PS1 cells following treatment with indicated siRNA. Bold points are coloured per biological replicate, with statistics by unpaired, two-tailed t-test \**P*<0.05, \*\*\**P*<0.001 (N=2, N=3). (E) Representative brightfield images of day 3 spheroids containing H2B-RFP MIA PaCa-2 cells and H2B-GFP PS1 cells, with cell type specific knockdown of either EPHRINB2 (MIA PaCa-2) or EPHB2 (PS1). Scale bar = 100 μm.

**Supplementary Figure 2.**
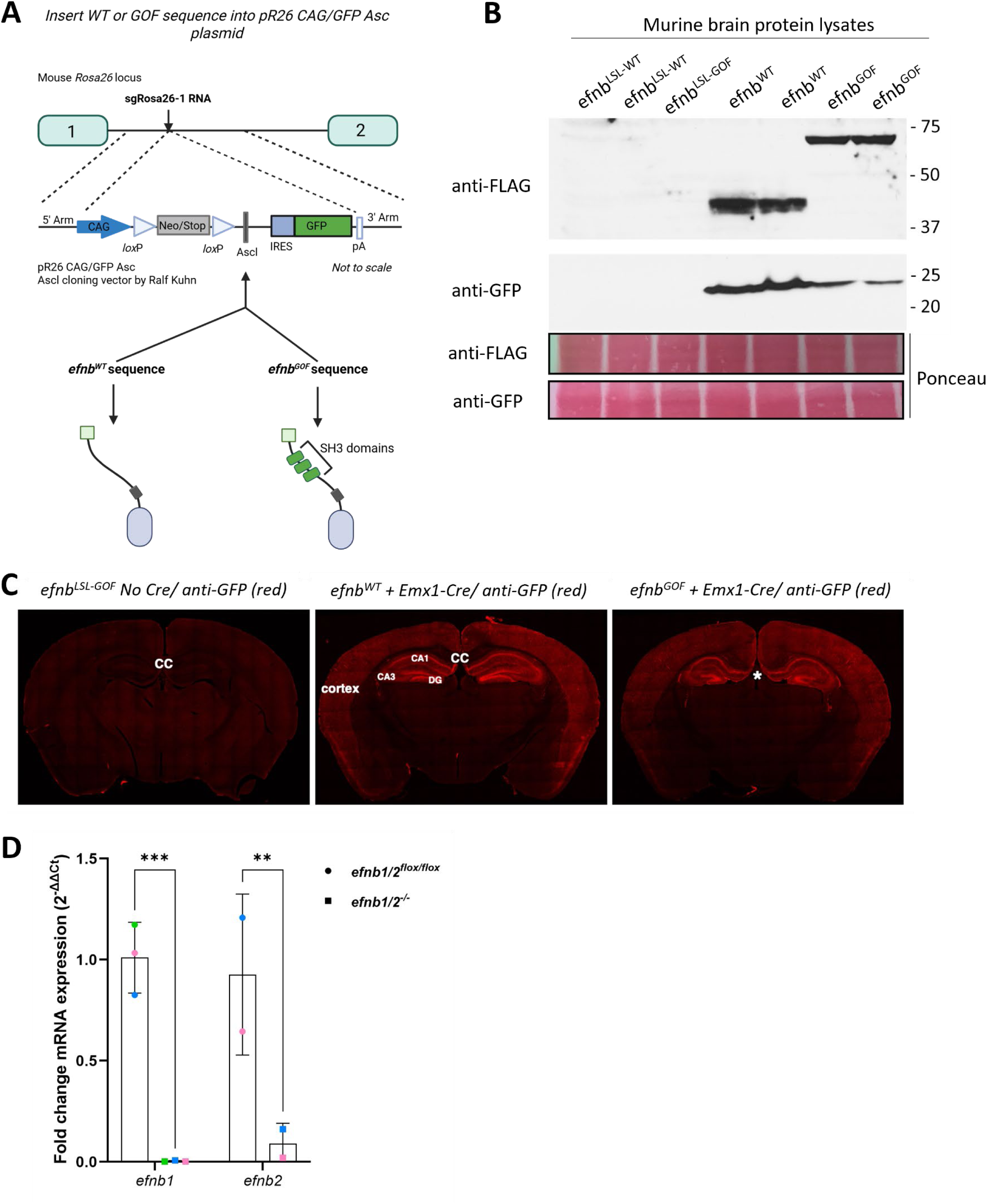
(A) Schematic of construct design for the *efnb^WT^* and *efnb^GOF^* cells, utilisng the pR26 CAG/GFP Asc cloning vector by Ralf Kuhn^48^. (B) Expression of Flag-tag and GFP protein from mouse brains with Rosa26 engineered WT and GOF that also contain Emx1-Cre driver. WT (40 kDa) and GOF (70 kDa) Ephrin protein expression is indicated. Ponceau staining highlights consistent sample loading. (C) GFP immunofluorescence (red) demonstrates restricted expression of GFP in cortex and hippocampus only in efnb^WT^ and efnb^GOF^ mice that also contained the Emx1-Cre driver. (D) qPCR quantification for *efnb1* and *efnb2* expression, comparing *efnb1/2^-/-^* to *efnb1/2^flox/flox^* stellate cells. Biological replicates are shown in different colours with two-way ANOVA and uncorrected Fisher’s least significant difference test used \*\**P*<0.01, \*\*\**P*<0.001 (N=3).

**Supplementary Figure 3.**
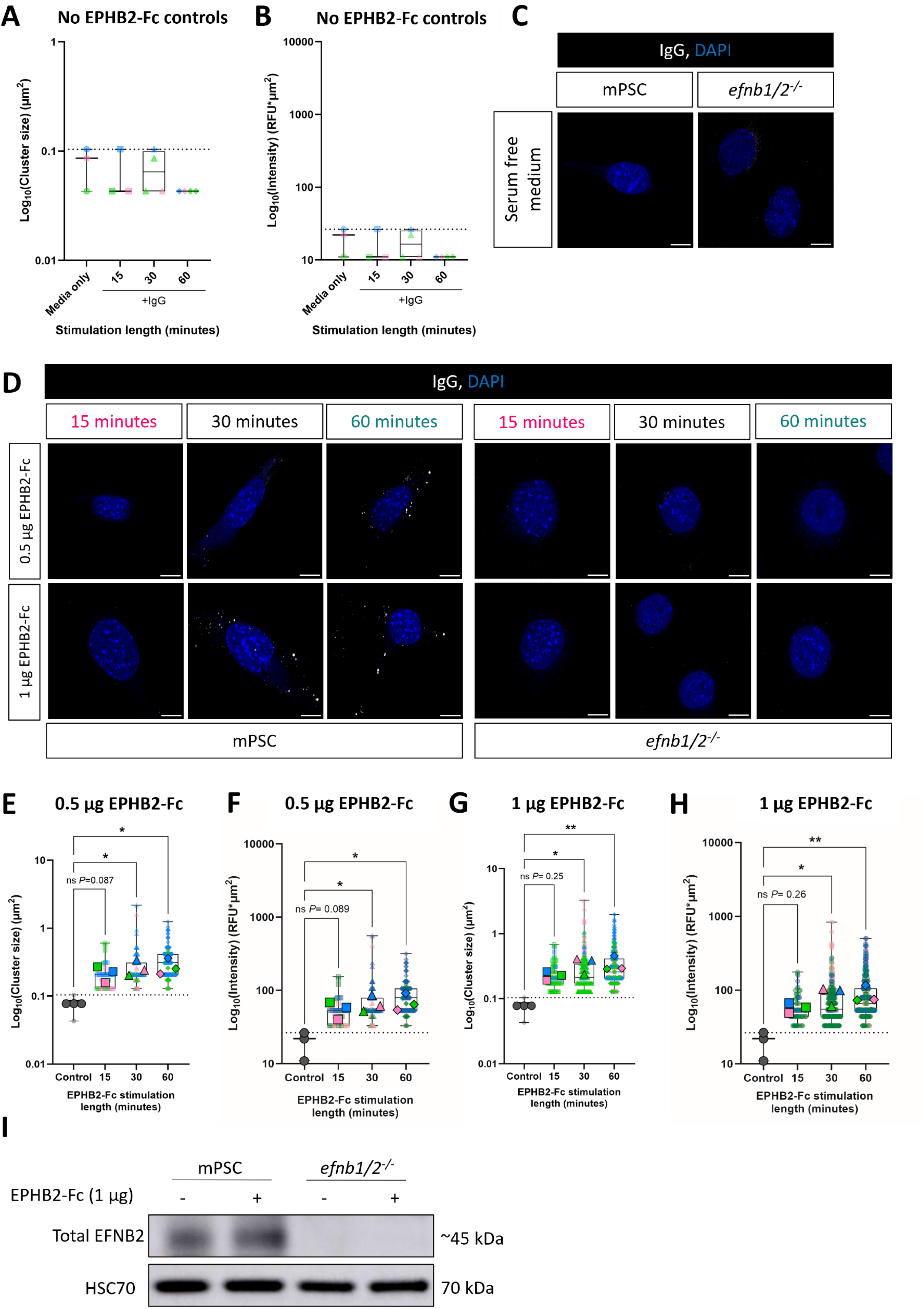
(A, B) Quantification of cluster size (A) and cluster intensity (B) for control measures which included IgG only or serum-free medium alone. Fiji ImageJ *analyse particles* function was utilised to quantify images. 1 point = 1 foci, with biological replicates shown in different colours (N=3). (C, D) Representative 63X confocal images of cluster formation in mPSC CAF or *efnb1/2^-/-^*stellate cells either untreated (C), or stimulated with either EPHB2-Fc (0.5, 1 μg) or IgG only (D) for 15, 30 or 60 minutes (N=3). Scale bar = 10 μm. (E-H) Quantification of cluster size (E, G) or cluster intensity (F, H) for 0.5 μg and 1 μg EPHB2-Fc stimulation. The control measures utilised (IgG only and serum-free medium alone) allowed background signal to be assessed (A-C). Only experimental values above this were plotted (dotted line). Superplots are shown with 1 point = 1 cluster, coloured by biological replicate with averages per repeat shown outlined and in bold. Kruskal Wallis with uncorrected Dunn’s test (E-H) \**P*<0.05, \*\**P*<0.01 (N=3). (I) Western blot of total EphrinB2 expression in CAF and *efnb1/2^-/-^* stellate cells. Both cell types were stimulated for 30 minutes with 1 μg EPHB2-Fc. HSC70 was used as a loading control (N=3).

**Supplementary Figure 4.**
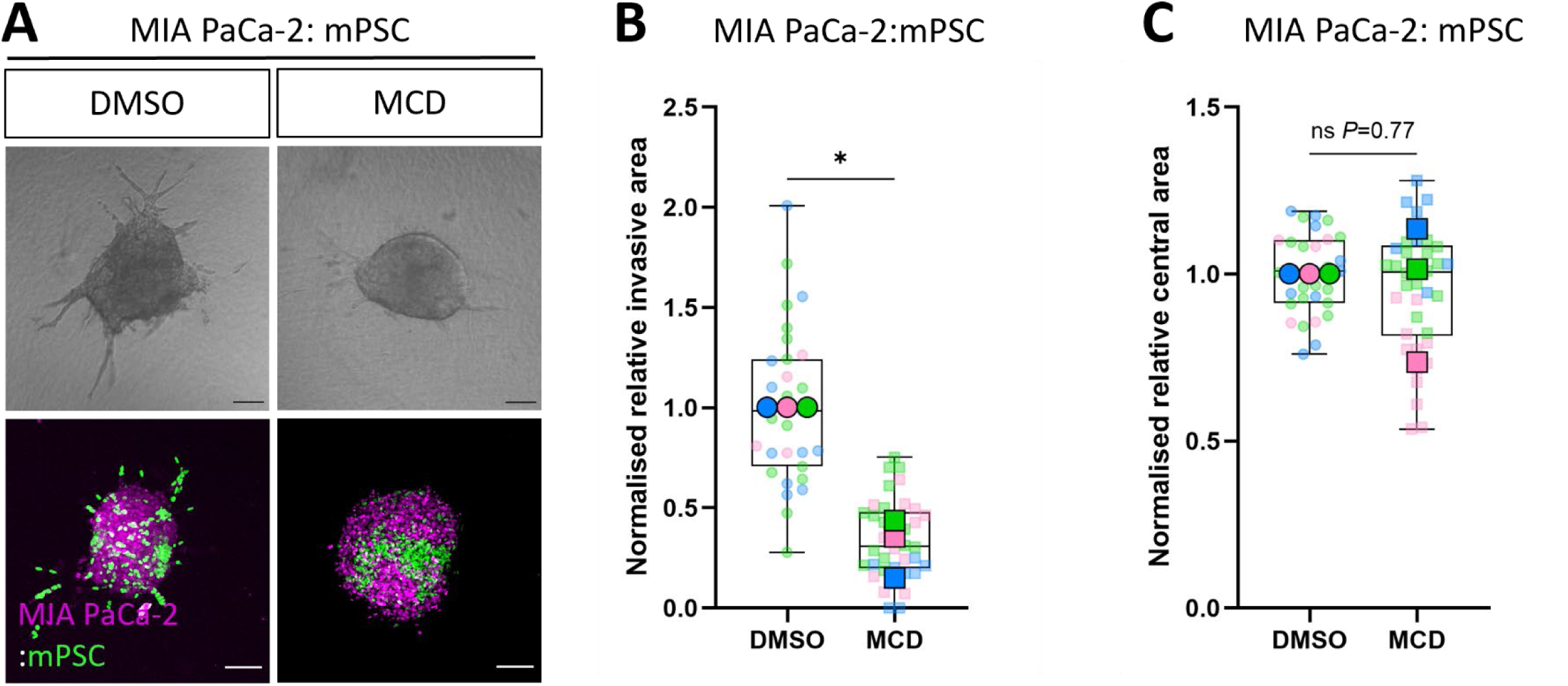
(A) Representative brightfield and confocal spheroid images of H2B-RFP MIA PaCa-2 (purple) and H2B-GFP CAF (green) cells. Spheroids were cultured for 3 days prior to imaging, with replenishment of medium containing DMSO or MCD (1 μM) every 24 hours. Scale bar = 100 μM. (N=3). (B, C) Spheroid quantification of invasive (B) and central (C) area comparing DMSO to MCD (1 μM). Graphs presented as superplots with 1 point = 1 cluster, coloured by biological replicate with averages per repeat shown outlined and in bold. Welch’s t-test utilised \**P*<0.05 (N=3).

**Supplementary Figure 5.**
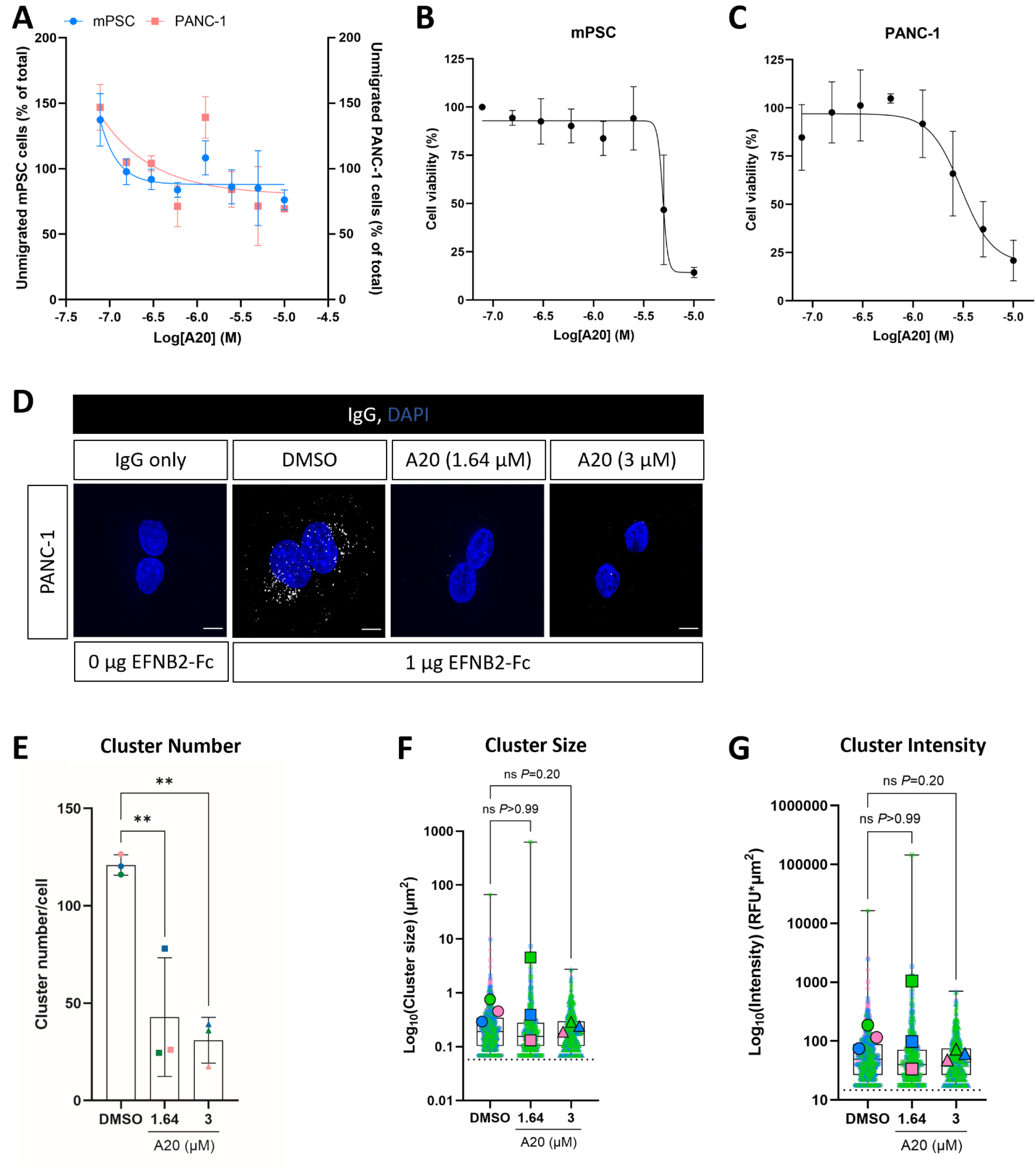
(A) Quantification of unmigrated cells in the apical compartment for PANC-1 (red) and mPSC CAF (blue) cells at varying concentrations of A20. Data presented as a percentage of the total number of cells initially plated (N=3). (B, C) MTT viability assay for mPSC (B) and PANC-1 (C) cells at varying concentrations of A-20 (N=3). (D) Representative 63X confocal images of cluster formation in PANC-1 PDAC cells following stimulation with and without 1 μg EFNB2-Fc for 30 minutes +/- indicated concentrations of A20 (blue indicates nuclei). Scale bar = 10 μm. (E-G) Cluster number per cell (E), cluster size (F) and cluster intensity (G) were quantified using the *analyse particles* macro on Fiji ImageJ. IgG alone values represented by dotted line. Superplots show all data above threshold (dotted line), with 1 point = 1 cluster, coloured by biological replicate with averages per repeat shown outlined and in bold. Ordinary one-way ANOVA with Dunnett’s multiple comparisons test (E), Kruskal Wallis with Dunn’s multiple comparisons test (F, G) utilised \*\**P*<0.01 (N=3).

**Supplementary Figure 6.**
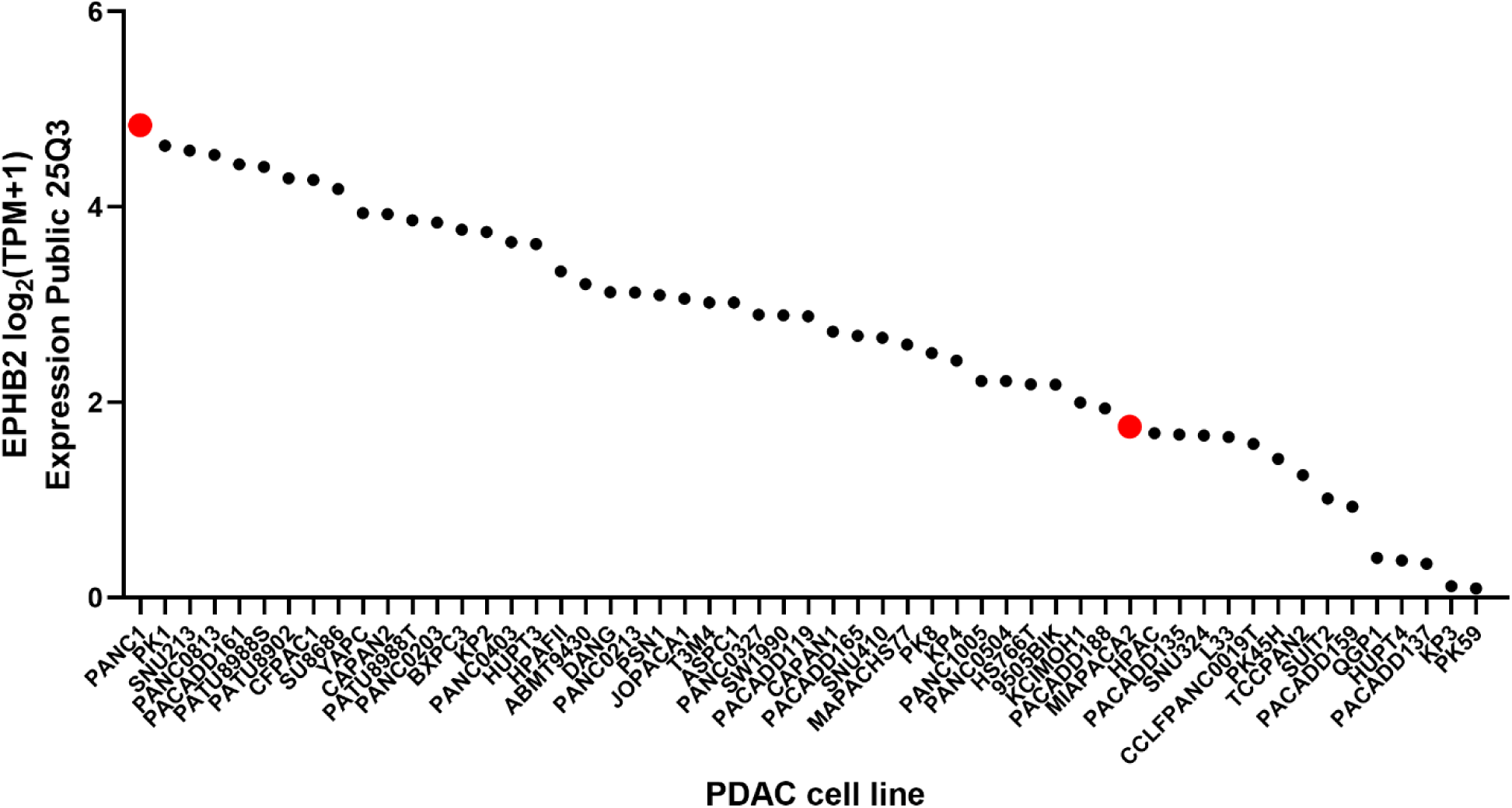
EPHB2 expression status of PDAC cell lines. Expression Public 25Q3 dataset was used from DepMap to plot EPHB2 expression across available PDAC cell lines. A high (PANC-1) and low (MIA PaCa-2) EPHB2-expressing cell lines are indicated in red.

**Supplementary Figure 7.**
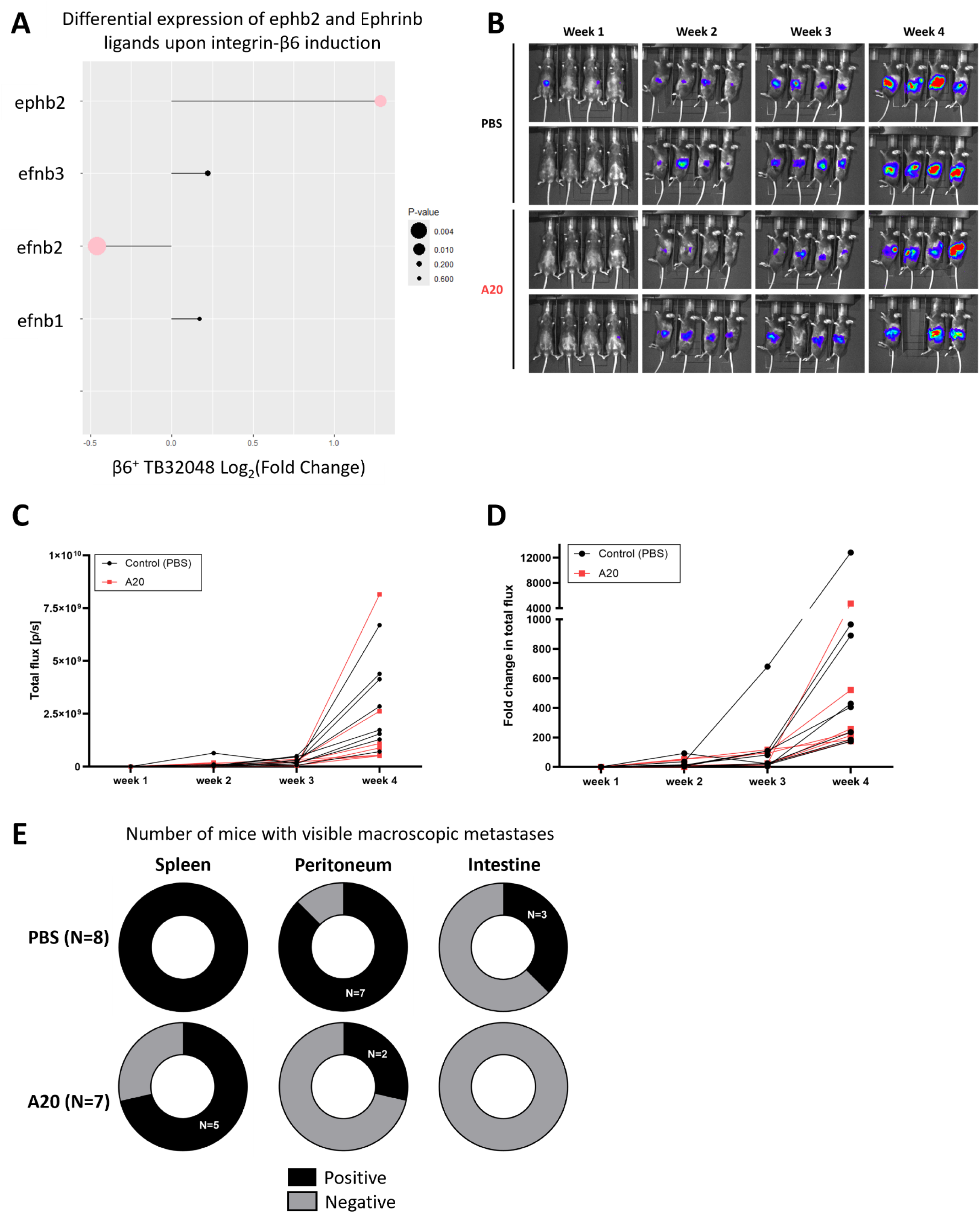
(A) Lollipop plot showing RNA-seq data from β6^+^ TB32048 tumours, highlighting a significant increase in ephb2 expression but a significant decrease in EphrinB2 expression. The size of the circle indicates the *P*-value, with significant values (*P*<0.05) shown in pink. (B) IVIS imaging of each mouse showing tumour formation across the 4-week experiment. (C, D) IVIS quantification showing total flux (C) and fold change in flux (D). Each mouse is shown separately with PBS control mice in black and A20 treated mice in red (N=8 PBS, N=7 A20). (E) Number of visible macroscopic metastases for spleen, peritoneum and intestine harvested from PBS (N=8) and A20 (N=7) treated mice. These measures were not counted as spontaneous metastases due to the proximity of each organ to the site of orthotopic injection.

**Supplementary Table 1.**
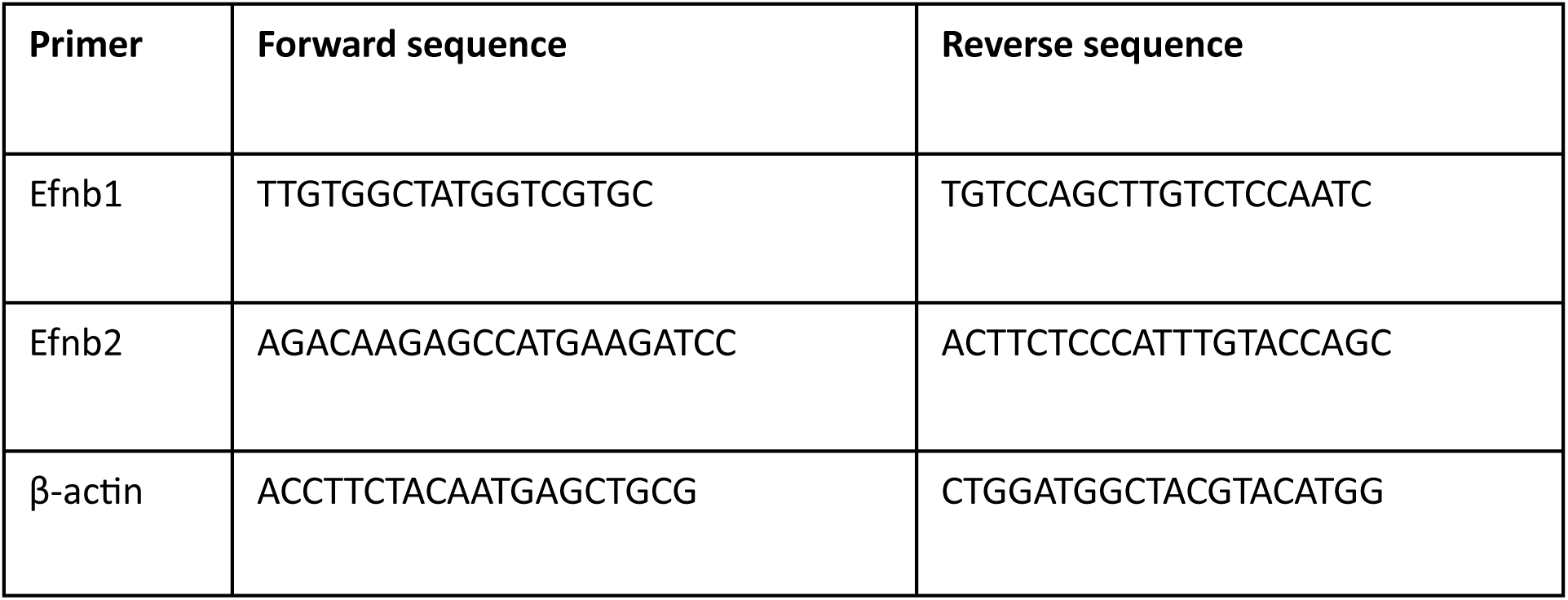
Primer sequences utilised in this manuscript.

